# Salivary proteins of *Phloeomyzus passerinii*, a plant-manipulating aphid, and their impact on early gene responses of susceptible and resistant poplar genotypes

**DOI:** 10.1101/504613

**Authors:** Luis Portillo Lemus, Jessy Tricard, Jérôme Duclercq, Quentin Coulette, David Giron, Christophe Hano, Elisabeth Huguet, Frédéric Lamblin, Anas Cherqui, Aurélien Sallé

## Abstract

Successful plant colonization by parasites requires the circumvention of host defenses, and sometimes a reprogramming of host metabolism, mediated by effector molecules delivered into the host. Using transcriptomic and enzymatic approaches, we characterized salivary glands and saliva of *Phloeomyzus passerinii*, an aphid exhibiting an atypical feeding strategy. Plant genes responses to salivary extracts of *P. passerinii* and *Myzus persicae* were assessed with poplar protoplasts of a susceptible and a resistant genotype, and in a heterologous *Arabidopsis* system. We predict that *P. passerinii* secretes a highly peculiar saliva containing effectors potentially interfering with host defenses, biotic stress signaling and plant metabolism, notably phosphatidylinositol phosphate kinases which seemed specific to *P. passerinii*. Gene expression profiles indicated that salivary extracts of *M. persicae* markedly affected host defenses and biotic stress signaling, while salivary extracts of *P. passerinii* induced only weak responses. The effector-triggered susceptibility was characterized by downregulations of genes involved in cytokinin signaling and auxin homeostasis. This suggests that *P. passerinii* induces an intracellular accumulation of auxin in susceptible host genotypes, which is supported by histochemical assays in *Arabidopsis*. This might in turn affect biotic stress signaling and contribute to host tissue manipulation by the aphid.

## 1 Introduction

In nature, plants engage in multiple interactions with a variety of organisms, resulting in either beneficial or detrimental impacts on plant performance. These interplays are mediated by effectors delivered into the host-plant, either in the apoplast or within plant cells [1, 2]. Effectors can be defined as molecules secreted by plant-associated organisms that alter host cell structure and function, positively or negatively depending on the partners involved [1]. This can be achieved through a direct modification of host molecules, or of their activity, by effectors with enzymatic or binding activities, or a modification of host gene expression by effectors with a nucleic acid binding activity [3].

Within the frame of host-parasite interactions, effectors have evolved to target various host functions and key processes. They primarily aim at suppressing plant immunity [1, 3]. For instance, parasites can target and inactivate plant second messengers involved in stress signaling such as Ca^2+^, reactive oxygen species (ROS) and extracellular ATP with cation-binding effectors, peroxiredoxins and apyrases, respectively [4, 5]. In some situations, effectors can also alter plant development, and promote the manipulation of plant resources by parasites [1, 3, 6]. For example, several pathogens secrete effectors which induce the production of sugar efflux transporters and hijack host sugars for their own nutrition [7]. Similar reconfiguration of host plant metabolism and tissues, to turn them into optimal substrates for development and fitness, have also been reported in plant pests [e.g. 6, 8, 9]. Successful interactions among parasite effectors and their targets lead to an effector-triggered susceptibility [3]. Nonetheless, resistant plant genotypes have evolved receptors, mostly with nucleotide-binding leucine-rich repeat (NB-LRR), that specifically interact with parasite effectors to induce an effector-triggered immunity [10]. Microbial pathogen effectors have been extensively studied [e.g. 11, 12]. Comparatively, effectors of insect herbivores and their impact on host plants have received only a limited attention until recently [13, 14]. Evidence indicates that both effector-triggered susceptibility and immunity also occur in plant-insect interactions [e.g. 15, 16]. Consequently, as for pathogens, a deep mechanistic understanding of effector action, i.e. an identification of effectors involved in the interplay and of their targets in host-plants, would be necessary to develop a sustainable management of agricultural and forestry pests, based on resistant plant genotypes [17].

The woolly poplar aphid, *Phloeomyzus passerinii* Sign., is a major pest of poplar plantations in the Mediterranean Basin, the Near East and France [18]. This aphid does not feed on sap but in the cortical parenchyma of its host-trees, where it induces a reaction tissue presumably acting as a physiological sink, draining nutrients from surrounding tissues [18, 19, 20]. During aphid outbreaks, when poplar trunks are covered with aphid colonies, the accumulation of reaction tissues dramatically affects host allocations of non-structural carbohydrates and nitrogen compounds, and infested mature trees can die rapidly, within a few months [18, 21]. Such outbreaks only occur in stands of susceptible poplar genotypes. Detailed histochemical investigations during early stages of infestation have unraveled that tissues affected by the unusually damaging probing activity of *P. passerinii* in these susceptible genotypes exhibit weak defense reactions, with limited lignification and accumulation of phenolic compounds [19, 20]. During later stages, a reaction tissue differentiate in the cortical parenchyma, characterized by an intense cell multiplication and hypertrophy, an accumulation of free and protein-bound amino acids, and an absence of both vacuolar phenolic compounds and starch granules [20, 22]. The differentiation of this reaction tissue improves both aphid performance and feeding behavior [23]. Conversely, in resistant poplar genotypes, an intense lignification and a marked accumulation of phenolic compounds occur readily after the onset of probing, blocking further differentiation of reaction tissues and preventing aphid colonization [18, 20]. It is therefore likely that *P. passerinii* induces either an effector-triggered susceptibility in susceptible poplar genotypes, allowing a beneficial manipulation of host tissue for the aphid, or a rapid and intense immunity response in resistant poplar genotypes preventing aphid nutrition. In-depth functional approaches would allow to unravel the molecular mechanisms that contribute to either the success or the failure of this pest establishment on its host-plant, which could ultimately help to improve poplar breeding programmes.

We hypothesize (i) that since it has a peculiar feeding strategy for an aphid [23], and it can hijack and reconfigure the metabolism of its host plant to improve its nutritional value, *P. passerinii* should possess a rich and atypical repertoire of salivary effectors compared to other sap-feeding aphids. Considering that susceptible hosts implement weak defense reactions when infested, we further hypothesize (ii) that salivary effectors successfully alter biotic stress signaling and / or defense molecules in these host genotypes to induce an effector-triggered susceptibility. Alternatively, in resistant hosts, the effector-triggered immunity should lead to a marked upregulation of defense-related genes. Finally, since it can manipulate host metabolism and anatomy, our last hypothesis (iii) is that some of its salivary effectors of *P. passerinii* can target genes involved in the cell-division cycle and / or induce the disruption or diversion of auxin and/or cytokinin-dependent pathways, which are important regulators of plant growth, differentiation and defense [6, 24].

More specifically, our first objective was to characterize with transcriptomic and enzymatic approaches the salivary gland encoded proteins and saliva, respectively, of *P. passerinii*. Our second objective was to assess *in vivo*, with a RT-qPCR approach, early responses of poplar genes belonging to different metabolic and signaling pathways using protoplasts exposed to aphid saliva. Protoplasts were used to simulate a host cellular response to salivary effectors of aphids. Our last objective was to investigate the impact of salivary extracts on specific gene expression *in planta*, in a heterologous *Arabidopsis* system. For all these steps comparative approaches were used. The predicted salivary gland encoded proteins of *P. passerinii* were compared with those of a sap-feeding aphid, *Myzus persicae* (Sulzer) [25, 26], in order to identify proteins common to both aphid species as well as proteins specific to *P. passerinii*. To get insight into how *P. passerinii* affects host immune system and metabolism during both an effector-triggered susceptibility and an immune response, the impact of salivary extracts on gene expression were assessed with poplar genotypes either susceptible or resistant to *P. passerinii*. To compare with non-host interactions, the protoplasts of these poplar genotypes were also exposed to salivary extracts of *M. persicae*, which do not feed on poplar.

## 2 Materials and Methods

### 2.1 **Plant and insect material**

Two aphid species were considered in our study: *P. passerinii*, our model species, and the green peach aphid, *M. persicae*. Both aphid species are Aphididae, but belong to different subfamilies (*P. passerinii* is the only member of the Phloeomyzinae while *M. persicae* is an Aphidinae), which diverged at 137 Ma [27]. Compared to *P. passerinii*, *M. persicae* exhibit a quite different feeding strategy since it is a sap-feeder, which host range does not include poplars, and preliminary establishment attempts confirmed that it cannot settle and develop on poplars (data not shown). Firstly, considering both species allowed to compare secretomes from two aphids with distinct feeding strategies, and to test whether the atypical feeding strategy of *P. passerinii* results in an unusual saliva composition, particularly enriched with potential effectors. Secondly, it allowed to compare poplar response during an interaction with a specialist parasite, to poplar response during a non-host species interaction, i.e. an interaction that would act as a positive control for defense reaction.

All individuals of *P. passerinii* used either for transcriptomic analyses of salivary glands or saliva collection originated from the same monoclonal colony established from an apterous parthenogenetic female, collected in 2013 in Brézé (France, 47°16’36’’ N 0°06’84’’ W, WGS-84). The colony was maintained in the laboratory on potted stem cuttings of I-214, a *Populus x canadensis* Moench. genotype, under 20 ± 1 °C, 70 ± 10% relative humidity and 16/8 h light/dark cycles. Individuals of *M. persicae* originated from a monoclonal colony established from an apterous parthenogenetic female collected in 1999 on a potato plant in Loos-en-Gohelle (France, 50°27’44” N 2°46’45” E, WGS-84). The colony was maintained under the same controlled conditions as *P. passerinii*, on turnips (Vilmorin).

Two *P. x canadensis* genotypes commonly planted in France, I-214 and Koster, were used for the experiments. *Populus x canadensis* is a host species for *P. passerinii*. However, both genotypes differ in their susceptibility to *P. passerinii*. I-214 is highly susceptible. The aphid shows high performance when fed with this genotype, and can induce its reaction tissue within its cortical parenchyma. Conversely *P. passerinii* cannot settle on Koster, and consequently induce a reaction tissue. This genotype is consequently considered to be highly resistant [18]. Considering these two poplar genotypes allowed to investigate interactions with *P. passerinii* leading either to an effector-triggered susceptibility, with the susceptible I-214 genotype, or to resistance, with the resistant Koster genotype. *Populus x canadensis* is not a host species for *M. persicae* and non-host species interactions between this aphid and both poplar genotypes were consequently expected. Stem cuttings (ca. 25 cm long, 2 cm diameter) were provided by the experimental nursery of Guéméné-Penfao (Office National des Forêts, France). They were collected in the autumn of 2016, and kept at 2°C, in dry conditions until use. In January 2017, the stem-cuttings were removed from storage and planted in 0.4 L pots, filled with a sterile sand-compost (50:50) mixture (Klasmann substrate 4 no. 267). The cuttings were then transferred to a growth chamber (20 ± 1°C, 70 ± 10% relative humidity, 16/8 h light/dark photoperiod, 2.65 kLx, and watered three times a week). Leaves were then used for protoplast production (see 2.3.3).

For *in planta* functional validation of the effects of salivary proteins, two transgenic lines of *Arabidopsis thaliana* (L.) Heynh. were used. The transgenic line *pIAA2*::GUS [28] was used to assess the auxin response, with its auxin-responsive promoter, while the cytokinin response was evaluated with *pARR16*::GUS, with its cytokinin-responsive promoter [29]. Seeds were sterilized with chloral gas, sown in Petri dishes on 0.8% (w/v) agar with 1% (w/v) sucrose-containing 0.5 Murashige and Skoog medium (MS), stored for 2 days at 4°C, and grown on vertically oriented plates in growth chambers under a 16/8 h light/dark photoperiod at 18°C.

## 2.2 Salivary transcriptome

### 2.2.1 Sample collection, RNA isolation and *de novo* transcriptome assembly

About 500 adults of apterous parthenogenetic *P. passerinii* aphids, collected directly on their host-plant, were dissected to collect pairs of salivary glands. Total RNA was extracted using the GeneJET RNA Purification kit (Thermo Fischer Scientific), according to manufacturer’s instructions. RNA was DNase treated using RNase-Free DNase Set (Qiagen). RNA concentration was measured using the Qubit® RNA Assay Kit (Life Technologies) and a Qubit® 2.0 Fluorometer (Invitrogen). Construction of cDNA-library and sequencing were performed by Eurofins® Genomics using a MiSeq v3 Reagent Kit (600 Cycles PE, Illumina, USA) and a MiSeq sequencer (Illumina), with 12.5 µg of total RNA. For the *de novo* transcriptome assembly, 15,453,942 pair-ended reads were sequenced and assembled using Velvet (v1.2.10; [30]) and Oases (v0.2.8; [31]) software tools (table S1). A multi-kmer approach was applied. Separate assemblies with different kmer lengths were conducted and the individual assemblies were merged to a final assembly. Kmer lengths of 69, 89, 109 and 129 were used. The separate assemblies were merged following the filter1-CD-HIT-EST procedure proposed in Yang & Smith [32]. A completeness assessment was performed using gVolante (v1.2.1; [33]) with the pipeline BUSCO v2/v3 and the reference dataset Arthropoda (table S1). This Transcriptome Shotgun Assembly project was deposited at DDBJ/EMBL/GenBank under the accession GHDF00000000. The version described in this paper is the first version, GHDF01000000.

### 2.2.2 Annotation, secreted proteins detection and identification

To perform comparisons with *M. persicae* the transcriptome of this aphid was retrieved on NCBI (http://www.ncbi.nlm.nih.gov/genbank/, accession numbers: <u>DW010205</u> - <u>DW015017</u>, <u>EC387039</u> - <u>EC390992</u>, <u>EE570018</u> - <u>EE572264</u>, <u>EE260858</u> - <u>EE265165</u>, <u>ES444641</u> - <u>ES444705</u>, <u>ES217505</u> - <u>ES226848</u>, and <u>ES449829</u> - <u>ES451794</u>). Salivary transcriptomes were annotated using the pipeline described in figure S1. Transcripts were first translated into amino acid sequences using Prodigal (v2.5; [34]). We then used the SignalP 4.0 Server (v4.1) to predict the presence of signal peptides and cleavage sites in the amino acid sequences [35]. To predict transmembrane domains, we submitted each amino acid sequence with a signal peptide to the TMHMM Server (v. 2.0; [36]). Putative proteins with a signal peptide and no transmembrane domain were considered to be potential secreted proteins. The sequences of complete ORFs without signal peptide were analyzed again with SecretomeP (v2.0; [37]). To remove mitochondrial proteins with a signal peptide, which are not secreted in the saliva, sequences were analyzed with TargetP (v1.1; [38]). Likewise, to remove proteins of the endoplasmic reticulum with a signal peptide, sequences were analyzed with PS-scan (Prosite pattern: PS00014), and with PredGPI [39] for glycosylphosphatidylinositol-anchor signals.

The remaining proteins were first mapped against the non-redundant protein sequences (nr) using Blastp (v2.3.0, NCBI, accessed on 03/30/2016), with an E-value cutoff at 1^e-3^. Protein domains were annotated with Blast2Go (v3.3; [40]), and InterProScan (v5.30-69; [41]). Whenever possible, protein sequences were assigned to Gene ontology (GO) terms with an E-value cutoff at 1^e-6^, enzyme codes (EC) and KEGG pathways.

OrthoVenn (http://aegilops.wheat.ucdavis.edu/OrthoVenn; [42]) was used to identify orthologous proteins within and between salivary transcriptomes of the two aphids. Intraspecific orthologous proteins are first grouped into clusters, which are then compared between species. Each cluster was annotated with the Uniprot database (http://www.uniprot.org; [43]) and the nr peptide sequence database (NCBI, accessed on 03/30/2016).

To detect proteins orthologous to salivary effectors of aphids, protein sequences of known aphid effectors, i.e. C002, ACE1, ACE2, ACYPI39568, ACYPI00346, MpC002, Mp1, Mp2, Mp42, Mp55, Me10, Me23 [44, 45, 46], were compared to the salivary transcriptome of *P. passerinii* with Blastp (E-value ≤ 1^e-3^).

### 2.3 Functional validation assays

#### 2.3.1 Aphid saliva collection

Aphids secrete two types of saliva within their host-plants, liquid and solid saliva [4]. The solid saliva is secreted during probing. It hardens rapidly and forms a solid sheath encasing aphid stylets within the host-plant, while the liquid saliva is secreted within cells and sieve tubes [4]. Both types of saliva contain effectors [e.g. 4, 47], and were collected in our experiments. Because of the particular trophic substrates of *M. persicae* and *P. passerinii* (i.e. sap and cortical tissues, respectively), a special protocol was used for each species. The saliva of *P. passerinii* was collected after incubation of 30 to 40 individuals of 2^nd^ and 3^rd^ instars aphids on sachets of Parafilm^©^ membranes containing an artificial diet [48]. The artificial diets were constituted by a disc of 0.5% (w/v) agar completed with 150 µL of 15% (w/v) sucrose. The saliva of *M. persicae* was collected after incubation of 30 to 40 individuals, of 3^rd^ and 4^th^ instars, on artificial diet containing 120 μL of a 15% (w/v) sucrose as previously described by Cherqui & Tjallingii [48]. Aphids were deposited in a feeding chamber during 24 h at 20°C, 60 ± 1% relative humidity and a 16/8 h light/dark period with 2.65 kLx. Feeding chambers containing the artificial diets, incubated in the absence of aphids, were used as control samples.

For *P. passerinii*, after 24 h aphid salivation, artificial diet discs were collected and transferred into 100 µL of TE buffer (10 mM Tris, 1 mM EDTA, pH 8). The salivary proteins were released from the artificial diet according to Yang et al. [49], with slight modifications. The tubes containing artificial diet discs were frozen in liquid nitrogen for 1 min, immediately thawed at 70°C for 3 min and then centrifuged at 11,000 x *g* for 20 sec. To discard the excess of agar, salivary extracts were centrifuged in Sartorius tubes with filters of 0.22 µm. The supernatant containing salivary proteins of *P. passerinii* were collected, pooled and then stored at −20°C. For *M. persicae,* after 24 h salivation, aphid saliva was collected according to Harmel et al. [25]. The artificial diet is collected containing soluble saliva. The solid saliva was collected during the rinsing of each lower Parafilm membrane with TE buffer containing 0.1% (w/v) of Tween 20 (TE/Tween). The extracts were centrifuged at 10,000 x *g* for 15 min. The salivary proteins in the pellet were collected, pooled with the soluble saliva and then stored at −20°C.

The sample containing protein saliva extracts were concentrated using 2 mL Vivaspin^©^ tube (Sartorius) with 3kDa cut-off. The tubes were then centrifuged at 5,000 x *g* for 70 to 120 min according to sample volumes, and proteins adhering to membranes were recovered by 100 µL of TE/Tween buffer. Control samples were prepared with artificial diets from feeding chambers without aphids. The protein quantification was performed by measuring absorbance at 280 nm with the NanoDrop^©^ 1000 (ThermoScientific).

#### 2.3.2 Enzyme activities

Several enzyme substrates were added to the previously described artificial diets with or without agarose to detect enzymatic activities present in saliva excreted from the aphid. To visualize proteins in the salivary sheaths, the lower Parafilm^©^ membranes were stained by adding a drop of 0.01% (w/v) Coomassie blue in 10% (v/v) glycerol for 2 h. Dihydroxyphenylalanine (DOPA), 0.1% (w/v) was added to identify phenoloxidase activity (PO; catechol oxidase, EC 1.10.3.1). The enzymatic product, melanin, should stain salivary sheaths and halos around the sheaths. To detect peroxidase (EC 1.11.1.7) activity, artificial diets were immersed for some minutes in 0.1% (w/v) diaminobenzidine (DAB, Sigma) in 50 mM Tris (pH 7.5) containing 0.1% (v/v) H_2_O_2_ (Sigma). The enzymatic product should induce reddish staining of salivary sheaths and halos. For identification of pectinase activity, 0.1% (w/v) of pectin (Sigma) was added to the medium. After exposure to aphids, the gel was transferred for 3 h into a Petri dish containing 50 mM citrate-phosphate buffer, at pH 5.0 to detect pectin (methyl) esterase (PME, EC 3.1.1.11) and at pH 6.4 to detect polygalacturonase (PG, EC 3.1.1.15). The gel was then stained with a solution of 0.01% (w/v) ruthenium red (Sigma) for 1 h, and then washed several times with distilled water. At pH 6.4, red halos around the salivary sheaths indicate PME activity, while non-staining halos at pH 5 in the pink pectin indicate PG activity. Finally, for proteinase activity (EC 3.4.99), 0.5% (w/v) of gelatin (Sigma) was added to the medium. After exposure to aphids, the medium was incubated overnight in a solution of 50 mM Tris (pH 8) containing 100 mM NaCl and 10 mM CaCl_2_, then stained with Coomassie blue. An absence of blue staining shows proteinase activity. All observations of proteins and enzymatic activities were performed by light microscopy (Axioplan 2, Zeiss, Jena, Germany).

#### 2.3.3 Poplar protoplast preparations and treatments

We wanted to assess early poplar gene responses to aphid saliva, as interactions between elicitors and effectors of herbivores and plant receptors occur readily after the onset of probing or feeding [50]. Once deposited on their plant material, adults and nymphs of *P. passerinii* can wait for several hours before probing and delivering saliva into host tissues, inject probably a varying amount of saliva and presumably affect a limited number of host cells [19, 20]. Therefore, we used protoplasts as plant material in order to (i) have a sufficient amount of plant material exposed to treatments for the RT-qPCR experiments, (ii) have an identical exposure duration and intensity to salivary extracts among biological replicates, and (iii) avoid the effects of physical stimuli exerted by aphid stylets on plant cells. Mesophyll protoplasts of the two poplar genotypes were obtained from young leaves as described in Wu et al. [51]. Leaves were cut into 1–2 mm fine strips in 0.3 M sorbitol and 66.67 mM CaCl_2_ (pH 5.6) and lysed in an enzyme solution (0.6 M mannitol, 0.25% (w/v) cellulase Onozuka R-10, 0.05% (w/v) macerozyme R-10) in the dark for 16 h with gentle shaking (30 rpm) at room temperature. Protoplasts were collected by filtering the lysate suspension through a 70 μm cell strainer (Falcon®) and concentrated by spinning down at ≈ 800 x *g* for 10 min at 4 °C. The pellet was washed twice with W5 buffer (154 mM NaCl, 125 mM CaCl_2_, 5 mM KCl, 5 mM glucose, 0.03% (w/v) MES, pH 5.8) and then resuspended in 0.6 M mannitol to a final concentration of 1 × 10^6^ protoplasts per mL. Protoplasts (1.10^6^) were incubated at 20°C with gentle shaking (40 rpm) for 3 h with aphid salivary proteins or with protein extraction buffer (control). This incubation duration allowed to observe early gene response to salivary effectors before cell walls start to replenish [52]. RNA extraction was done just after the saliva or control treatment (see 2.3.4). Preliminary experiments investigating expression of 5 poplar genes, and conducted with 1, 10, 20, 40 or 80 µg of salivary proteins, indicated that the optimal response (i.e. the maximum fold change) was observed with 1 µg of salivary proteins of *P. passerinii* and 10 µg of salivary proteins of *M. persicae* (Fig. S2). Protoplast viability, before and after treatment with aphid saliva, was assessed using 0.005% (w/v) fluorescein diacetate (FDA). After 5 min of incubation protoplasts were observed under blue light epifluorescence, and cell viability was estimated as the percentage of fluorescent cells. Most protoplasts were intact and viable after enzymatic digestion with cellulase and macerozyme (98%), as well as after incubation with salivary proteins (95%; Fig. S3).

#### 2.3.4 Quantitative RT-PCR

After the aphid saliva treatments, protoplasts were centrifuged at ≈ 800 x *g* for 2 min. Total RNAs were then immediately extracted with the RNeasy® Plant Kit Mini Kit (Qiagen). A DNase treatment with the RNase-free DNase Set (Qiagen) was carried out for 15 min at 25°C. Total RNA concentration was determined using a Nanodrop ND-1000 spectrophotometer. All RNA samples were rejected if they did not reach a minimum concentration of 100 ng μL^−1^, a 260 nm/280 nm ratio between 1.8 and 2.0. Poly(dT) cDNA was prepared from 1 μg total RNA using the iScriptTMcDNA Synthesis Kit (Bio-Rad) and quantified with a LightCycler 480 (Roche) and SYBR GREEN I Master (Roche), according to the manufacturer’s instructions. PCR was carried out in 384-well optical reaction plates heated for 10 min to 95°C to activate hot start Taq DNA polymerase, followed by 40 cycles of denaturation for 60 sec at 95°C and annealing/extension for 60 sec at 58°C. The distribution of the quantitative RT-PCR mix containing SYBR Green I Master (Roche), cDNAs and primers was performed using the EVO150^©^ (Tecan) pipetting robot in a 384-well plate. The expression of 43 genes, belonging to eight different physiological processes or metabolic pathways (i.e. auxin, cytokinins, jasmonates, ethylene, salicylic acid, phenolic compounds, reactive oxygen species (ROS), cell-division cycle), was quantified with specific primer pairs designed by Quant-Prime [53] based on the *Populus trichocharpa* sequence (v3.0) from Phytozome (https://phytozome.jgi.doe.gov/pz/portal.html; Table S2). Expression levels were normalized to the levels of *PtUBIQUITIN10* (*PtUBQ10*), commonly used as a reference gene in plants [e.g. 54]. All RT-qPCR experiments were done with three independent biological replicates, with two technical replicates each. One of the biological replicates of the *M. persicae* – Koster interaction was excluded from the analyses because of technical issues during quantification. Relative gene expression was calculated according to the ^ΔΔ^Ct method, with protoplasts incubated with protein extraction buffer as controls. Primers used for gene expression analysis are listed in Table S2.

#### 2.3.5 Histochemical analysis of GUS activity

Transgenic seedlings of *A. thaliana* (five-day-old and eight-day-old for *pIAA2*::GUS and *pARR16*::GUS, respectively) were incubated with 2 mL of liquid MS containing 1 µg and 10 µg of aphid salivary proteins (in TE/Tween buffer) of either *P. passerinii* or *M. persicae* for 3 h and 4 h for *pIAA2*::GUS and *pARR16*::GUS, respectively. Positive controls were incubated with 20 µM of indole acetic acid (IAA) (Sigma-Aldrich), and 20 µM of 6-benzylaminopurine (BAP) (Sigma-Aldrich). Negative controls were incubated in liquid MS and corresponding volumes of TE/Tween buffer. Five seedlings were used for each modality. Seedlings were then incubated in reaction buffer containing 0.1 M sodium phosphate buffer (pH 7), 2 mM ferricyanide, 2 mM ferrocyanide, 0.1% (v/v) Triton X-100 and 1 mg ml^−1^ X-Gluc for 1 up to 24 h in dark at 37 °C. Afterwards, chlorophyll was removed by destaining in 70% ethanol and seedlings were cleared as described by Malamy and Benfey [55]. GUS expression was monitored by differential interference contrast microscopy.

#### 2.4 Data analysis

All tests were carried out with the statistical software R 2.11.0 [56]. RT-qPCR results were expressed as fold-changes in gene expression compared to the reference gene *PtUBQ10*. Fold-changes were log_2_ - transformed. Following this transformation, fold-changes varied between -∞ and +∞, with negative values corresponding to gene underexpression, positive values to overexpression and zero, to no change in gene expression. To visualize similarities and differences in fold-changes a heatmap was built, with genes in columns and modalities in lines. The heatmap was built with a Z-score, i.e. log_2_ – transformed fold changes which were normalized and centered per column. Hierarchical clustering, using Pearson correlation as similarity metric, was performed to visualize the proximity among genes in columns and among modalities in lines. The uncertainty of hierarchical clustering of modalities was assessed with approximately unbiased p-values calculated with a multiscale bootstrap resampling, with 10,000 bootstrap replications, using the function pvclust() [57]. T-tests were performed with ^Δ^Ct values to detect significant changes in expression for each saliva treatment vs. controls. An analysis of variance, followed with a Tukey test when significant, was also performed to compare the effect of treatment modalities (i.e. aphid species and host genotype combinations) on log_2_-fold changes of each gene.

## 3 Results

### 3.1 **Annotation, secreted proteins detection and identification**

From 36,312 and 3,233 transcripts, 1,243 and 221 transcripts were predicted to encode for secreted salivary proteins in *P. passerinii* and *M. persicae*, respectively. About half of them (604) were annotated for *P. passerinii* and 190 for *M. persicae*. Using OrthoVenn, 121 and 58 protein clusters were identified for *P. passerinii* and *M. persicae*, respectively. About 17% of these clusters were common between the two aphids (Table S3).

Blast2GO determined that *P. passerinii* salivary gland encoded proteins were predominantly binding proteins and enzymes (Fig. 1, table S4). The most common enzymes were peptidases (especially serine-type and cysteine-type endopeptidases), kinases (especially phosphatidylinositol phosphate (PIP) kinases) and hydrolases. Several enzymes involved in the degradation of carbohydrates (i.e. cellulase, trehalase, β-glucuronidase, mannosidase and glucosylceramidase), and of phenolic compounds (i.e. peroxidase and oxidoreductase) were also identified (Fig. 1, table S4). Among binding proteins, dimerization protein, nucleic acids binding (especially DNA binding), nucleotide binding (especially ATP binding) and cation-binding (mostly calcium ion-binding and zinc-binding proteins) were the most commonly found (Fig. 1, table S4). Proteins related to hormone activity were also identified. Glucose dehydrogenases were also detected with OrthoVenn (table S3). Among the 12 aphid salivary effectors considered, five were identified in *P. passerinii*, with low E-values (< 7 e^-71^): Mp10, ARMET, ACE 1, ACE2 and ACE3.

**Figure 1:**
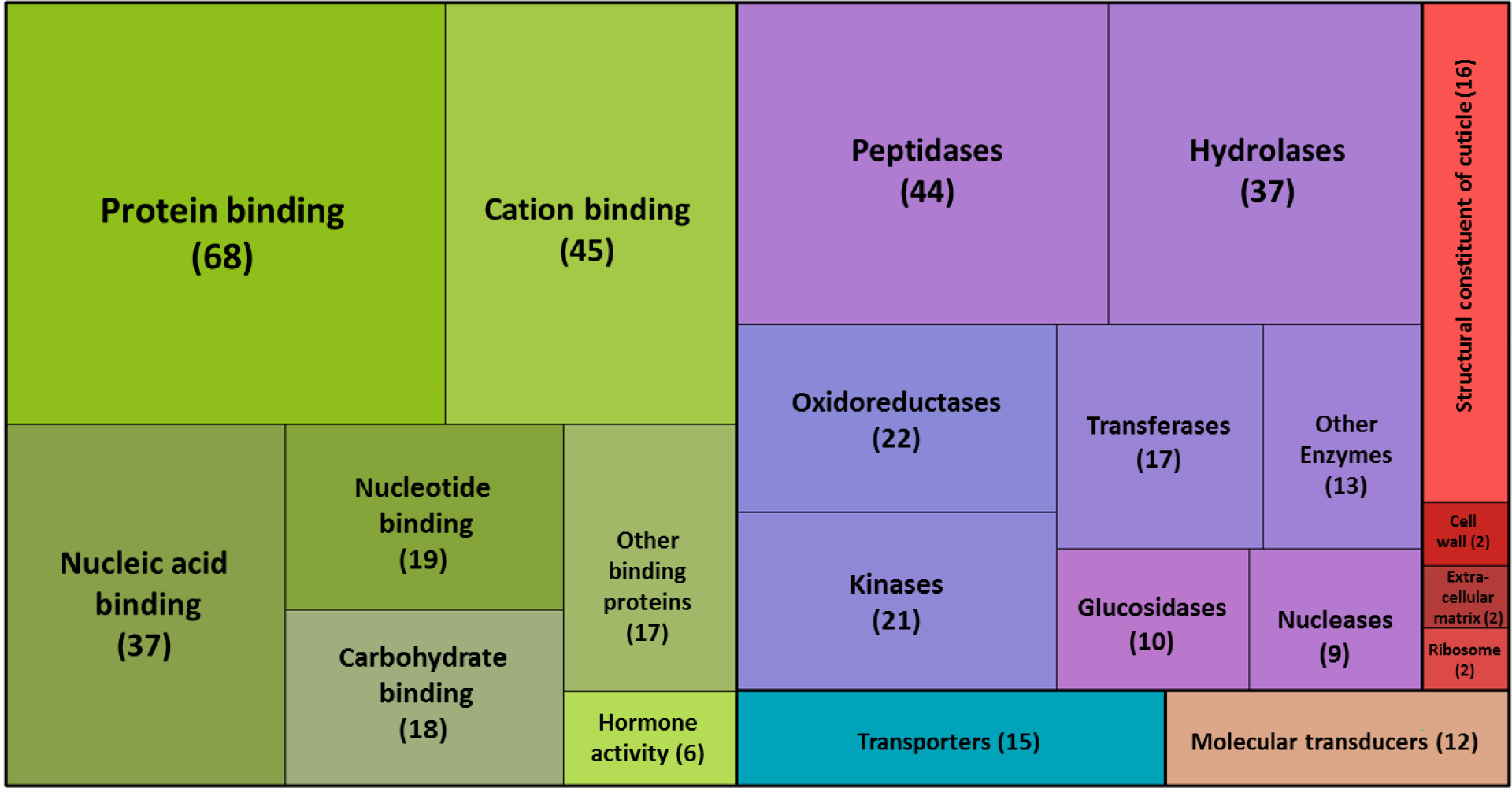
Gene Ontology treemap for the salivary transcriptome of *Phloeomyzus passerinii*. The box size correlates to the number of sequences isolated. Numbers between brackets indicate the number of sequences identified. Green boxes indicate binding proteins, purple boxes indicate enzymes, red boxes indicate structural constituents, the blue box indicates transporters and the brown box molecular transducers. The treemap was created with the treemap() function in R. A Detailed list of the proteins can be found in the table S4.

### 3.2 Enzyme activities

Staining with Coomassie blue confirmed the protein nature of the salivary sheath material (Fig. 2A) and DOPA staining indicated a phenoloxidase activity in the sheaths (Fig. 2B). Black halos were also observed around some sheaths (Fig. 2B). Likewise, peroxidase activity was found in salivary sheaths and halos around sheaths (Fig. 2C). However, no pectinesterase, polygalacturonase and proteinase activity was detected.

**Figure 2:**
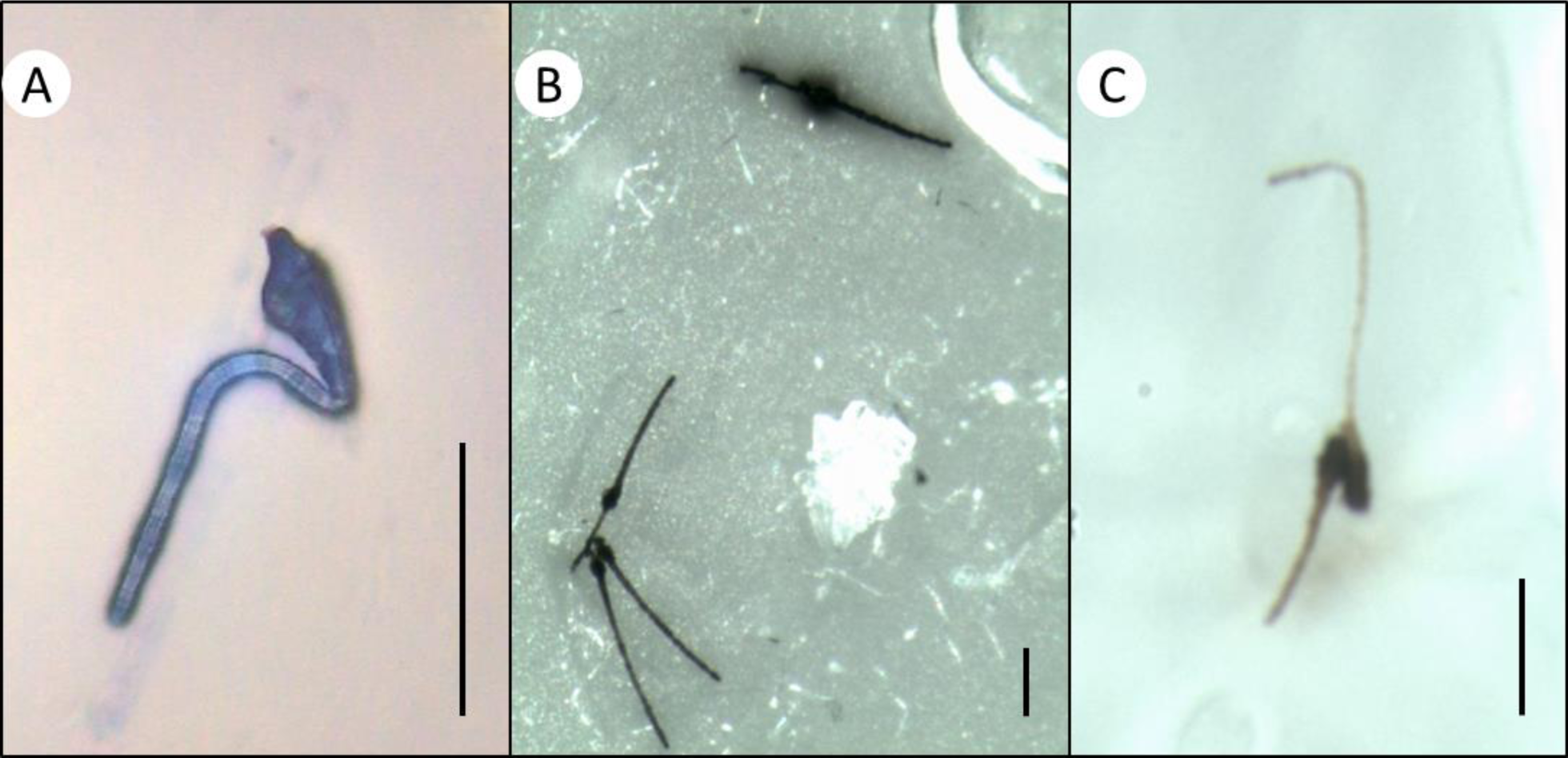
Representative salivary sheaths secreted in artificial diets by *Phloeomyzus passerinii*. Sheaths stained and observed after 24 h probing in an agarose diet: (A) sheath stained with Coomassie blue; (B) black stained sheaths in diet containing 0.1% DOPA, indicating a phenoloxidase activity, note the dark halo surrounding the upper sheath; (C) reddish stained sheath in diet immersed with 0.1% DAB and 0.1% H_2_O_2_, indicating a peroxidase activity. Black bars represent 10 μm.

### 3.3 Quantitative poplar gene expression

An overview of poplar genes responses to aphid saliva is presented in figure 3. The hierarchical clustering of modalities (left dendrogram) showed that all biological replicates of interactions with the salivary proteins of *M. persicae* (i.e. non-host species interactions) were grouped within a significant cluster, in which a majority of poplar genes were upregulated. Within this cluster, replicates were not arranged according to poplar genotype. Biological replicates of the interaction between salivary proteins of *P. passerinii* and Koster (i.e. the host species – resistant genotype) also clustered together, although the cluster was significant for only two replicates. In this group, a majority of genes were weakly expressed or upregulated. For biological replicates of the interaction between salivary proteins of *P. passerinii* and I-214 (i.e. the host species - susceptible genotype), a majority were weakly expressed or downregulated. Nonetheless, these replicates were not significantly clustered together. The hierarchical clustering of genes (upper dendrogram) showed that there was no clustering of genes according to physiological process or metabolic pathway.

**Figure 3:**
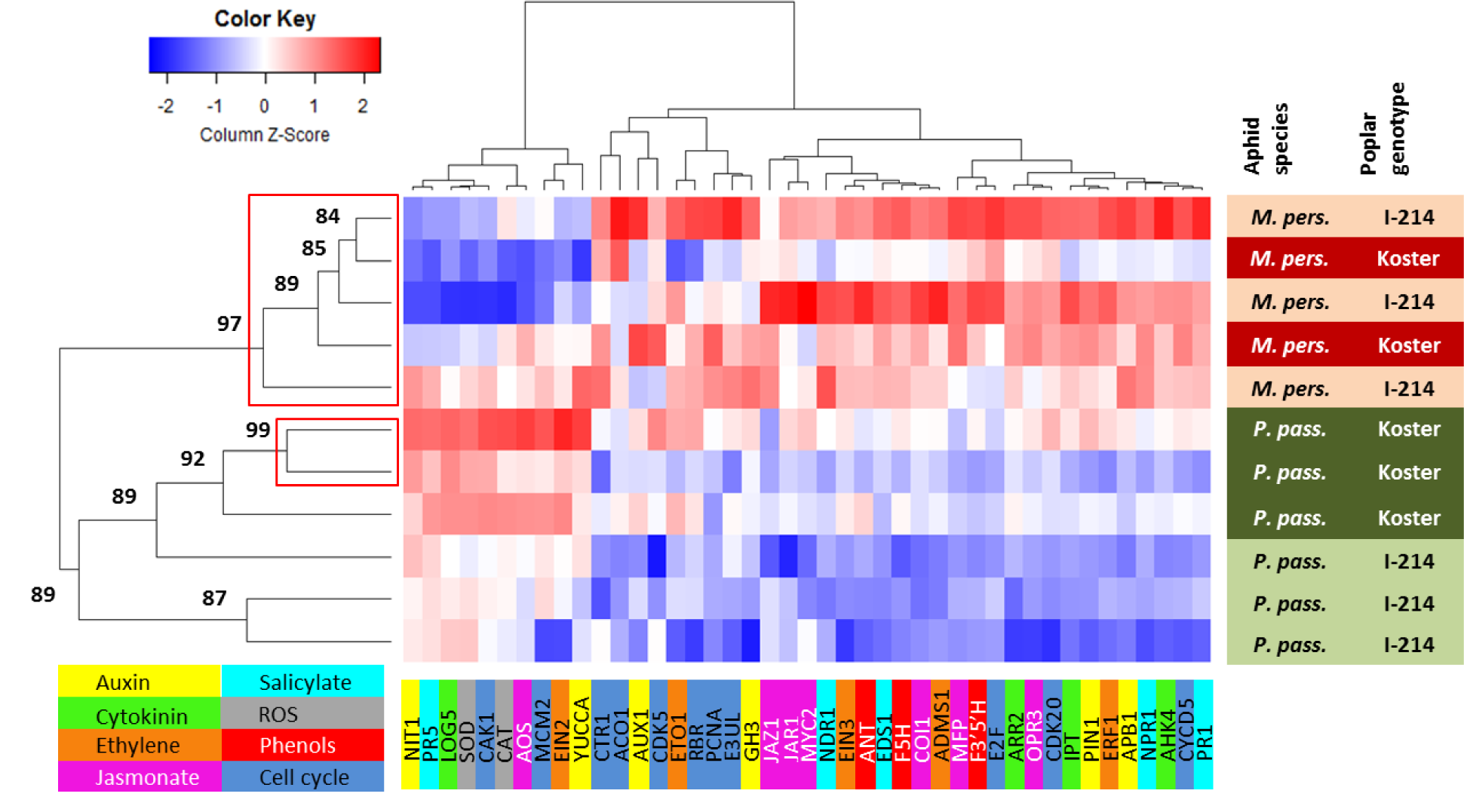
Heatmap of log_2_-fold changes of 43 poplar genes belonging to eight different physiological processes or metabolic pathways (lower left box), after incubation of poplar protoplasts of two poplar genotypes (Koster and I-214) with salivary proteins of two aphids (*Myzus persicae* (*M. pers.*) and *Phloeomyzus passerinii* (*P. pass.*)). Non-host species interactions are expected between both poplar genotypes and *M. persicae*. Poplar is a host plant for *P. passerinii* but Koster is a resistant genotype while I-214 is a susceptible genotype. Downregulation appears in blue and upregulation in red. Gene code is presented below the heatmap, modalities (i.e. aphid x poplar genotype combinations are presented on the right of the heatmap). Hierarchical clustering was built with distances based on Pearson correlations. Approximately unbiased p-values (%) are indicated for the hierarchical clustering of modalities, red boxes indicate significant clusters at α = 0.05.

For the auxin pathway, during non-host species interactions between salivary proteins of *M. persicae* and both poplar genotypes biosynthetic genes were either not affected (*PtYUCCA*) or downregulated (*PtNIT1*) while *PtGH3*, which is involved in auxin inactivaton via conjugation, was upregulated (Fig. 4). *PtAPB1* (signaling) and *PtPIN1* (auxin efflux carrier) were also significantly upregulated, but only during the interaction between *M. persicae* and I-214. No change was observed in the expression of the auxin influx carrier *PtAUX1*. For *P. passerinii*, during the interaction with Koster these genes displayed an expression profile quite similar to that observed during non-host species interactions with *M. persicae*, except for *PtNIT1* whose expression was not affected. The expression of *PtGH3* was significantly downregulated during the interaction between salivary proteins of *P. passerinii* and I-214, and *PtGH3*, *PtAPB1* and *PtPIN1* were differentially expressed during the interaction with this susceptible genotype compared to non-host species interactions with *M. persicae*, and compared to the interaction with the resistant genotype for *PtAPB1* only (Fig. 4).

**Figure 4:**
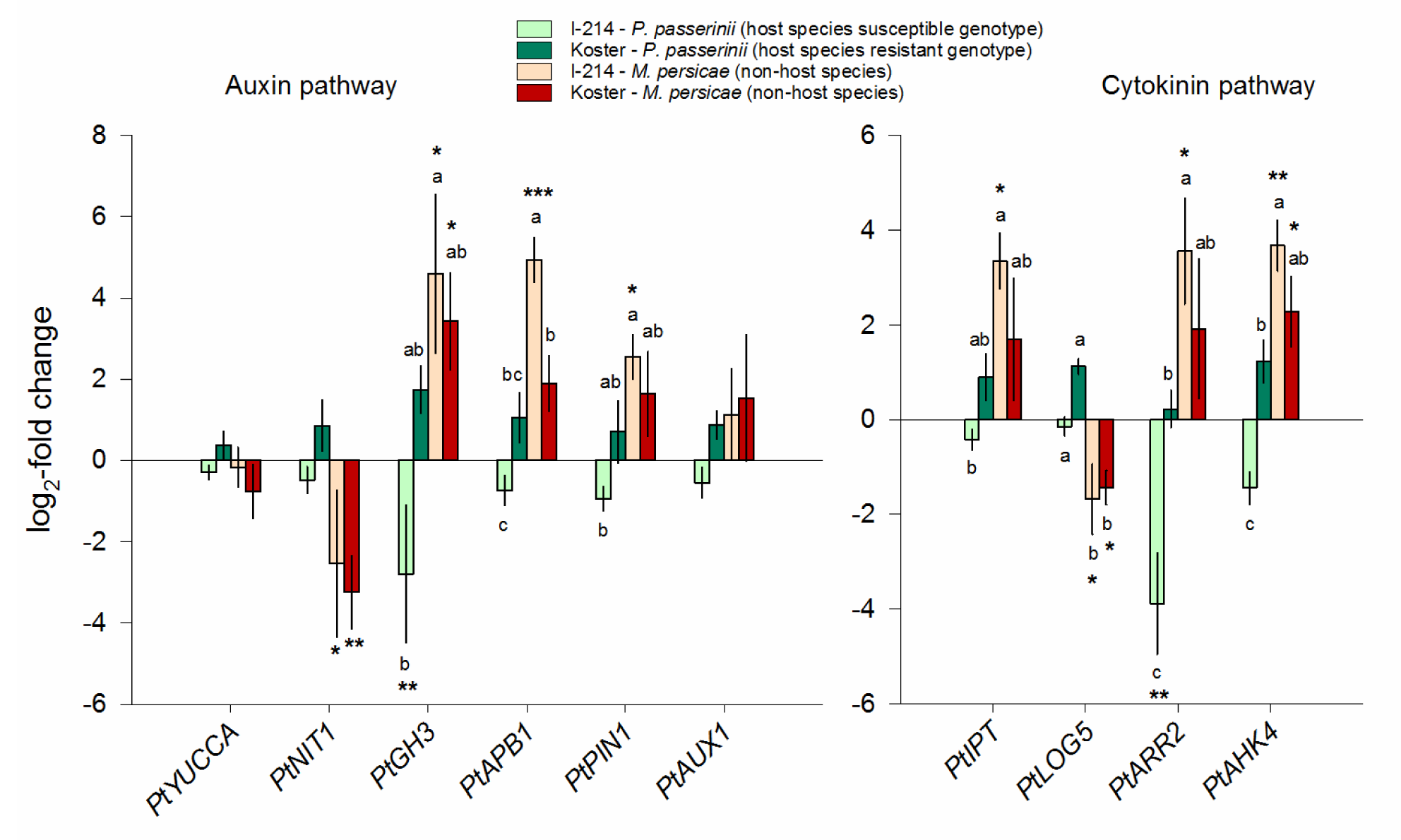
Mean (± SE) log_2_-fold changes of genes involved in auxin (left) and cytokinin (right) pathways of poplar protoplasts collected from two poplar genotypes (I-214 and Koster), after incubation with salivary proteins of two aphids (*Myzus persicae* and *Phloeomyzus passerinii*), resulting in resistant or susceptible interactions with a host species or non-host species interactions. Stars indicate modalities for which a significant upregulation or downregulation of gene expression vs. controls was observed, using ^Δ^Ct values (t-test). ***, ** and * indicate a significant effect with P < 0.001, P < 0.01 and P < 0.05, respectively. Different letters indicate significantly different groups (Tukey test) of log_2_-fold changes among modalities, for each gene.

For the cytokinin pathway, during the non-host species interaction between salivary proteins of *M. persicae* and I-214 *PtIPT* (biosynthesis), *PtAHK4* (perception) and *PtARR2* (signaling) were upregulated while *PtLOG5* (metabolism) was significantly downregulated (Fig. 4). A similar expression profile was observed during the non-host species interaction with Koster, but only *PtAHK4* and *PtLOG5* were significantly affected. These genes were not significantly affected during the interaction between salivary proteins of *P. passerinii* and Koster. In contrast, *PtARR2* was strongly repressed during the *P. passerinii* – I-214 interaction, and *PtAHK4* and *PtARR2* were differentially expressed during this interaction compared to the other ones (Fig. 4).

Regarding biotic stress signaling (i.e. jasmonates, salicylic acid, ethylene and ROS), non-host species interactions were characterized by a marked upregulation of a majority of genes, which was more frequently significant for the interaction with I-214 than for the interaction with Koster (Fig. 5). Few genes in jasmonate ethylene and ROS pathways were not affected by salivary extracts of *M. persicae* (i.e. *PtAOS*, *PtJAZ1*, *PtEIN2*, *PtETO1*, *PtCAT* and *PtSOD*), and, in the salicylic acid pathway, *PtPR5* was significantly downregulated. For interactions with salivary proteins of *P. passerinii*, all genes were weakly downregulated during the interaction with the susceptible genotype. In contrast, during the interaction with the resistant genotype most genes were either not affected or exhibited profiles as during non-host species interactions with *M. persicae*, with weak upregulations. Nonetheless, for these modalities, no gene expression significantly differed from the controls (Fig. 5). In most cases, gene expressions during the interaction with the susceptible genotype were significantly lower than during non-host species interactions with I-214, and, in fewer cases, lower than during non-host species interactions with Koster. For interactions with *P. passerinii*, only *PtCOI1* and *PtEIN3* genes were differentially expressed during interactions with either the susceptible or the resistant genotype. Conversely, since gene expressions during interactions with the resistant genotype were weakly upregulated, they generally did not differ from those observed during non-host species interactions with Koster, and differed from non-host species interactions with I-214 for only several genes (i.e. *PtCOI1*, *PtEDS1*, *PtNPR1*, *PtNDR1*, *PtEIN3*, *PtADMS1*). Nonetheless, *PtPR5* and *PtSOD* were clearly differentially expressed between host and non-host interactions with Koster.

**Figure 5:**
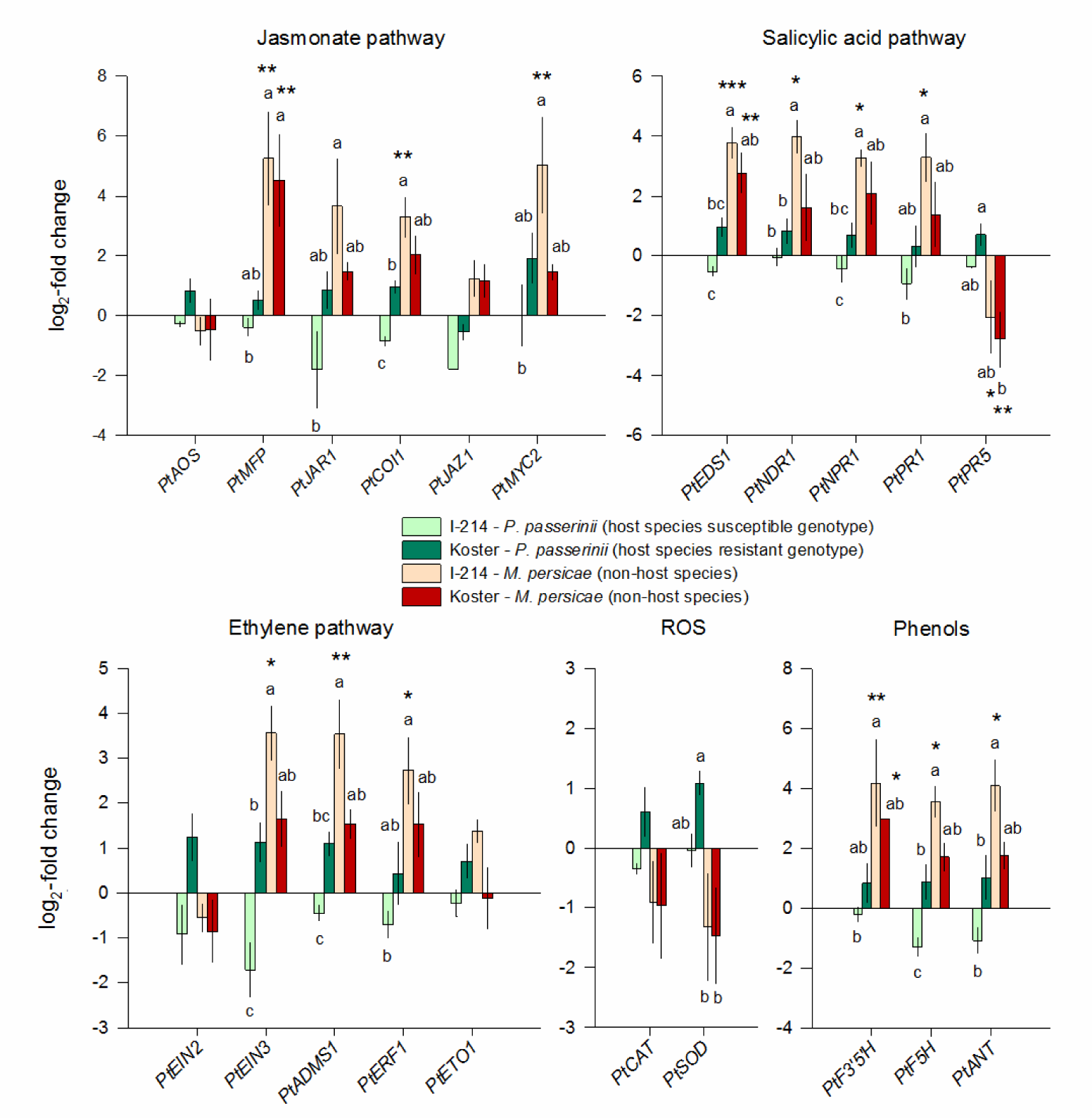
Mean (± SE) log_2_-fold changes of genes involved in jasmonate, salicylic acid, ethylene pathways, ROS responses and phenolic compounds of poplar protoplasts collected from two poplar genotypes (I-214 and Koster), after incubation with salivary proteins of two aphids (*Myzus persicae* and *Phloeomyzus passerinii*), resulting in resistant or susceptible interactions with a host species or non-host species interactions. Stars indicate modalities for which a significant upregulation or downregulation of gene expression vs. controls was observed, using ^Δ^Ct values (t-test). ***, ** and * indicate a significant effect with P < 0.001, P < 0.01 and P < 0.05, respectively. Different letters indicate significantly different groups (Tukey test) of log_2_-fold changes among modalities, for each gene.

For genes involved in the phenolic compounds pathway (Fig. 5), non-host species interactions, especially with I-214, led to an upregulation of all the considered genes. All genes were weakly, but not significantly, downregulated or upregulated during interactions between the saliva of *P. passerinii* and either the susceptible genotype or the resistant genotype, respectively. For I-214, gene expression was always significantly lower during the interaction with the saliva of *P. passerinii* compared to the interaction with the saliva of *M. persicae*. For *PtF5H*, expression was also significantly lower during the *P. passerinii* – I-214 interaction compared to other interactions.

Finally, regarding genes involved in cell-division cycle, *PtCDK5*, *PtPCNA*, *PtCDK20 and PtE2F* were significantly upregulated during non-host species interactions with *M. persicae*, while *PtCAK1* was significantly downregulated (Fig. 6). *PtCYCD5* and *PtACO1* were also upregulated but only during non-host species interactions with I-214. Likewise, *PtCDK5* and *PtCDK20* were also significantly upregulated during the *P. passerinii* - Koster interaction, as well as *PtCAK1*. Expression of *PtMCM2* and *PtRBR* was not affected by salivary extracts. *PtCDK20* was differentially expressed during the *P. passerinii* - I-214 interaction compared to other ones. Likewise, during the *P. passerinii* - Koster interaction the expression of *PtCTR1* was much lower compared to non-host species ones and expression of *PtE3UL*, *PtE2F* and *PtACO1* were lower compared to expression during non-host species interactions with I-214. For *PtCAK1*, the expression was higher during the *P. passerinii* - Koster interaction than during non-host species ones.

**Figure 6:**
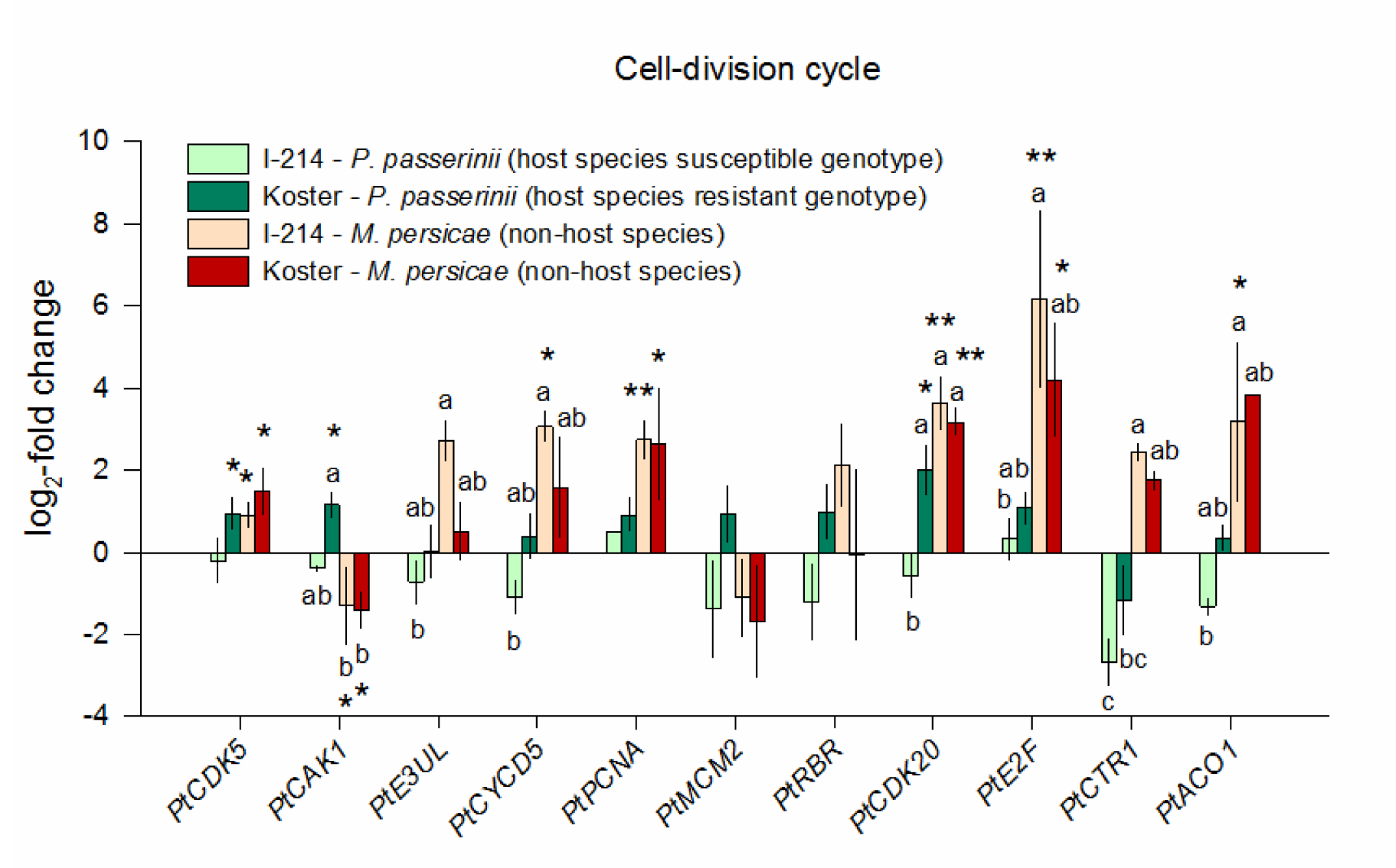
Mean (± SE) log_2_-fold changes of genes involved in cell-division cycle of poplar protoplasts collected from two poplar genotypes (I-214 and Koster), after incubation with salivary proteins of two aphids (*Myzus persicae* and *Phloeomyzus passerinii*), resulting in resistant or susceptible interactions with a host species or non-host species interactions. Stars indicate modalities for which a significant upregulation or downregulation of gene expression vs. controls was observed, using ^Δ^Ct values (t-test). ** and * indicate a significant effect with P < 0.01 and P < 0.05, respectively. Different letters indicate significantly different groups (Tukey test) of log_2_-fold changes among modalities, for each gene.

### 3.4 Histochemical analysis of GUS activity

Salivary extracts of *P. passerinii* increased *pIAA2*::GUS signals (Fig. 7E, 7F, 7I, 7J), which were similar to those caused by an exogenous application of auxin (Fig. 7A and 7B). Incubation with salivary proteins of *M. persicae* resulted in faint colorations (Fig. 7G, 7H, 7K, 7L), similar to those of negative controls (Fig. 7C and 7D).

**Figure 7:**
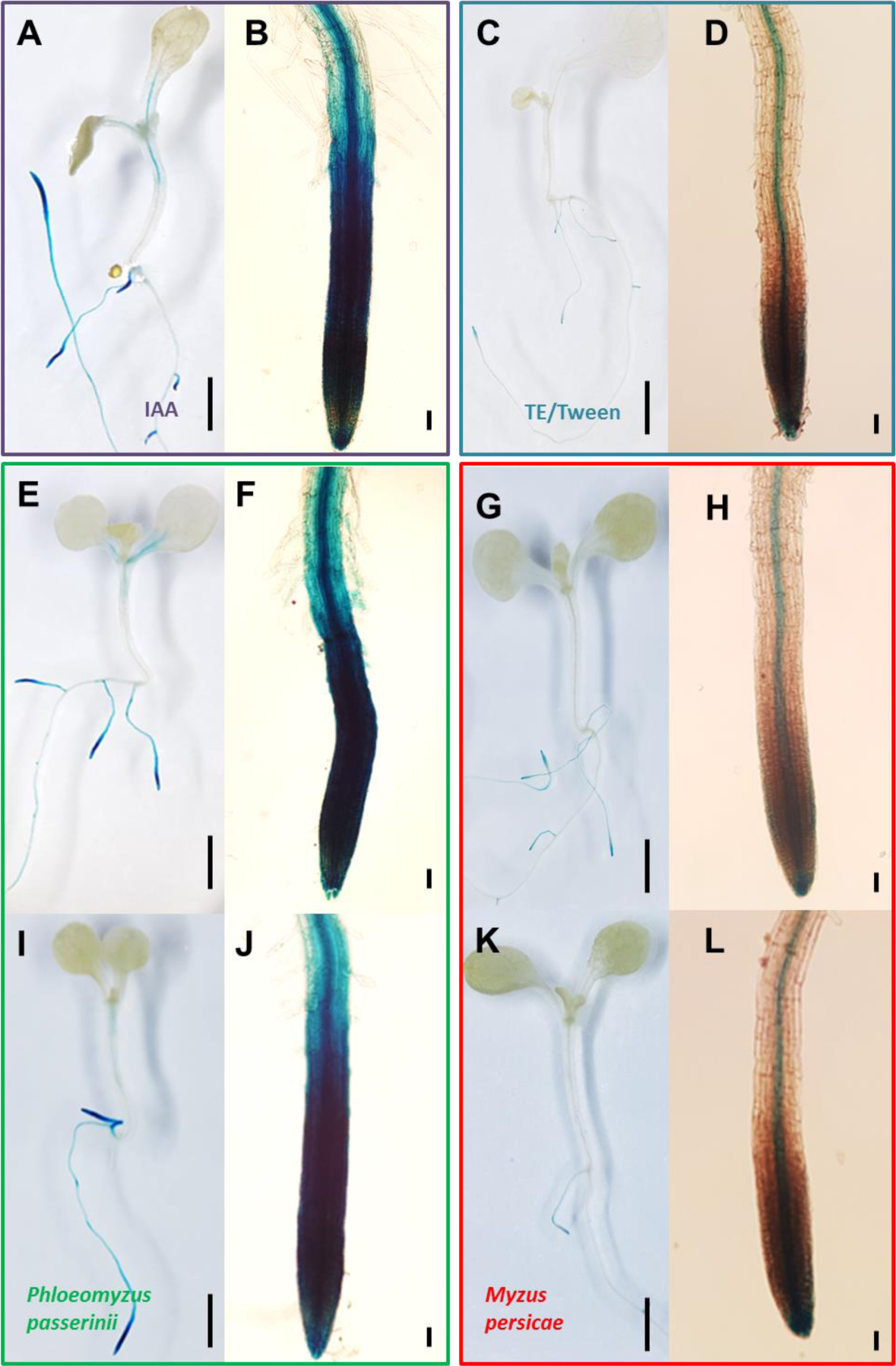
Representative GUS assays of transgenic seedlings of *Arabidopsis thaliana pIAA2*::GUS, showing whole plants (A, C, E, G, I and K), and root tips (B, D, F, H, J and L), after 3 h of incubation in 20 µM of IAA (A and B), TE/Tween buffer (C and D), and 1 and 10 µg of salivary proteins of *Phloeomyzus passerinii* (E, F, I and J) or *Myzus persicae* (G, H, K and L). Black bars represent 1 mm for whole plants (A, C, E, G, I and K) and 10 µm for root tips (B, D, F, H, J and L). Five seedlings were used for each modality.

Positive controls of *pARR16*::GUS were characterized by a strong staining in the middle part of root central cylinder (Fig. 8A and 8B), which was weak in negative controls as well as with *M. persicae* salivary proteins (Fig. 8G, 8H, 8K, and 8L). No coloration was visible in the roots of seedlings incubated with salivary proteins of *P. passerinii* (Fig. 8E, 8F, 8I and 8J).

**Figure 8:**
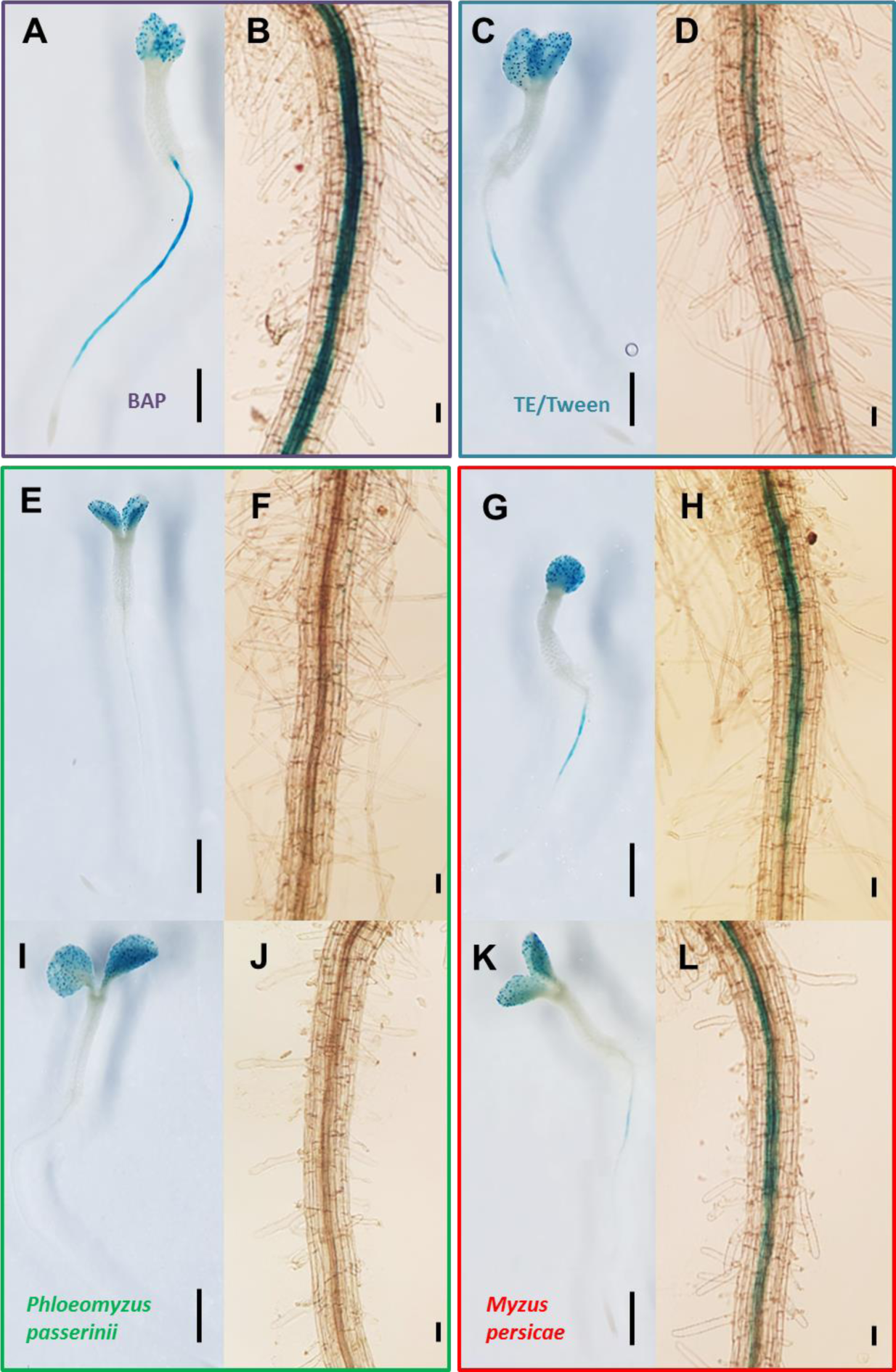
Representative GUS assays of transgenic seedlings of *Arabidopsis thaliana pARR16*::GUS, showing whole plants (A, C, E, G, I and K), and root tips (B, D, F, H, J and L), after 4 h of incubation in 20µM of BAP (A and B), TE/Tween buffer (C and D), and 1 and 10 µg of salivary proteins of *Phloeomyzus passerinii* (E, F, I and J) or *Myzus persicae* (G, H, K and L). Black bars represent 1 mm for whole plants (A, C, E, G, I and K) and 10 µm for root (B, D, F, H, J and L). Five seedlings were used for each modality.

## 4 Discussion

Successful plant colonization by parasites generally requires the circumvention or inactivation of host defenses, and sometimes a finely tuned reprogramming of host metabolism. As a consequence, the lifestyle and feeding strategy of parasites should have both quantitative and qualitative outcomes on their secretome [e.g. 11, 58]. In line with this, comparisons among salivary proteins of aphid species also indicated that aphids with different host species and / or feeding strategies exhibited very different salivary protein profiles [59, 60]. Our results tend to support this assertion since the predicted salivary gland proteins of *P. passerinii* and *M. persicae* shared few similarities, and poplar gene expression profiles markedly differed when protoplasts were exposed to salivary proteins of these two aphid species. The salivary transcriptome of *P. passerinii* also hold specific and abundant PIP kinases sequences, which have never been reported from any other aphid saliva. Nonetheless, several protein functions predicted in our study (e.g. calcium-binding, DNA-binding, ATP-binding, GTP-binding proteins, glucose dehydrogenases, oxidoreductases, trehalases and phosphatases) have also been identified in the saliva of different sap-feeding aphid species [22, 59, 61, 62], and in secretions of other plant parasites [3, 63]. Interestingly, calcium-binding proteins are supposedly key components of the saliva of sap-feeding aphids, preventing the plugging of sieve tubes [4]. Since *P. passerinii* does not feed on sap [19], it suggests that these proteins also play other crucial roles during aphid-plant interactions. In addition to their different feeding strategies, the two aphid species are also phylogenetically distant [27], which should further increase the dissimilarity of their secretomes. It should be noted also that the transcriptomes of *P. passerinii* and *M. persicae* have been obtained with a different sequencing and assembly methods [64]. This probably explains the difference in transcripts amounts gathered for both aphids and several encoded proteins in salivary glands of *M. persicae* might be missing in its transcriptome, leading to an apparently low similarity between secretomes. Likewise, the lack of biological replicates for both transcriptomes might have also led to an underestimation of the similarity between secretomes.

Several encoded proteins detected in the salivary glands of *P. passerinii* may interfere with host defenses *via* direct interactions with secondary metabolites or defense proteins. The interaction between *P. passerinii* and susceptible poplar genotypes is characterized by transient accumulation of phenolic compounds, which fade away during later stages of the interaction [20]. Salivary effectors might contribute to the degradation of these secondary metabolites as numerous oxidoreductase sequences were identified in the salivary gland transcriptome of *P. passerinii*. Moreover, *in situ* biochemical assays confirmed the presence of active peroxidases and phenoloxidases in both solid and soluble saliva fractions of *P. passerinii*. The predicted glucose dehydrogenases may also help *P. passerinii* to detoxify defensive compounds of the host-plant [11, 60]. Likewise, plant defense proteins might be inactivated by the numerous protein-binding proteins and proteases predicted among the salivary gland encoded proteins of *P. passerinii*. Parasites frequently secrete similar effectors to degrade or modulate the plant enzymes activities [3]. Nonetheless, no protease activity was observed during *in situ* bioassays. Gelatin was probably not the adequate substrate to detect the protease activity of *P. passerinii*. Additional *in situ* assays could be conducted to detect, in *P. passerinii* saliva, the activity of the proteases and possibly other enzymes like cellulases. A proteomic analysis of salivary extracts should also confirm and complement the predictions of our transcriptomic approach [65, 66].

Several salivary gland encoded proteins may also affect host defense *via* a disruption of biotic stress signaling. PIP kinases catalyze phosphorylation of phosphatidyl-inositol into phosphatidylinositol-4-5-biphosphate (PIP_2_). The hydrolysis of PIP_2_ produces secondary messengers like inositol-1-4-5-triphosphate (IP_3_) and diacylglycerol (DAG). This latter can in turn be hydrolyzed into phosphatidic acid (PA) considered an important signaling molecule in plants, triggered in response to various biotic and abiotic stresses [67]. PIP_2_ and IP_3_ might also affect cellular oscillations of cytosolic Ca^2+^ and are involved in multiple processes including cell cycle and phytohormone regulation [68]. Other proteins, also frequently detected in parasites, including aphids, may also interfere with secondary messengers like calcium-binding, ATP-binding, and GTP-binding proteins or with hormone signaling like hormone-binding proteins [3, 5, 6, 59]. Similarly, it has been hypothesized that trehalase may interfere with trehalose-based defense responses in *A. thaliana* [61].

Finally, several salivary gland encoded proteins could also contribute to the manipulation of host-plant metabolism. For instance, nucleic acid-binding proteins could affect gene expression, while protein-binding and hormone-binding proteins could modulate metabolic and phytohormonal pathways [3, 59]. In addition, serine proteases, acid phosphatases, cellulases, lipases and metalloproteases have also been predicted or detected in the secretions of different gall-inducing organisms, and supposedly contribute to gall induction and / or maintenance [63, 69].

Both the *in vivo* approach with protoplasts and the heterologous *in planta* assay with *A. thaliana* confirmed that salivary proteins of *P. passerinii* impact plant gene transcription, in a specific manner. Gene expression profiles markedly differed, especially in terms of intensity, between non-host species interactions with *M. persicae* and host species interactions with *P. passerinii*. Non-host species interactions led to an upregulation of most genes involved in jasmonate, ethylene and salicylic acid pathways, which are typically activated following aphid feeding, together or separately, depending on the aphid – plant interaction system considered [70, 71, 72]. As a consequence, genes related to secondary metabolism, i.e. *PtF3’5’H, PtANT* and *PtF5H*, were also upregulated during these interactions. Conversely, the expression of genes related to biotic stress signaling and defense was either unaffected or slightly downregulated or upregulated during interactions with a susceptible and a resistant genotype of the host species, respectively. This suggests that salivary effectors of *P. passerinii* manage to inactivate or bypass the biotic stress signaling in their host plant, as frequently observed in plant-parasites interactions [1, 3]. Non-host species interactions with the two poplar genotypes led to similar gene expression profiles while, for interactions with *P. passerinii*, the expression of several genes differed according to host genotype. Among them, the expression of several genes involved in biotic stress signaling (*PtCOI1*, *PtEIN3*), and defense (*PtF5H*), was significantly higher during the interaction with the resistant genotype than during the interaction with the susceptible one. The overall trend of lack of response or downregulation of genes involved in defense or biotic signaling observed during the interaction between salivary extracts of *P. passerinii* and the susceptible host genotype is congruent with gene expression profiles observed during effector-triggered susceptibility in other plant-parasite systems [73, 74].

Although different from the interaction with a susceptible genotype, the interaction between the resistant genotype and *P. passerinii* did not lead to marked upregulations of genes related to biotic stress signaling or defense. Gene expression profiles of poplar protoplasts during the interaction between *P. passerinii* and the resistant poplar genotype shared many similarities with those observed during non-host species interactions with *M. persicae*. Both types of interactions were characterized by an overall upregulation of host genes, which was generally more important during non-host species interactions than during host species interaction with the resistant host genotype, for which only few significant upregulations were observed. Interestingly, two of the three genes that were differentially expressed during non-host and incompatible interactions on the same poplar genotype (i.e. *PtPR5* and *PtSOD* – with a similar trend for *PtCAT*) were related to biotic stress signaling, and were slightly upregulated during the interaction with the resistant genotype while they were significantly downregulated during non-host species interactions. Effector-triggered immunity is generally a strong defense response, associated with extensive and rapid transcriptional reprogramming [10, 74, 75]. This was not detected during interactions between *P. passerinii* and the resistant host genotype. Nonetheless the reprograming can take place within few hours [75], and the incubation duration might have been too short to detect an extensive reprogramming. Slightly longer incubation durations might solve this issue.

To knock-down or divert stress signaling parasites may also reconfigure auxin and cytokinin signaling pathways [1, 6, 76]. Both RT-qPCR experiments and histochemical assays confirmed that *P. passerinii* can actively manipulate both auxin and cytokinin pathways during an interaction with a susceptible host genotype. Salivary extracts of *P. passerinii* did not affect either auxin biosynthesis or auxin transporter genes such as *PtPIN1* and *PtAUX1*, but significantly downregulated *PtGH3*, which is involved in the homeostasis of active auxin forms [77]. This could lead to intracellular accumulation of active auxin forms, which is supported by the activation of the auxin-responsive promoter *IAA2* during *in planta* assays with transgenic seedlings of *A. thaliana*. Auxin accumulation could in turn interfere with salicylic acid signaling and defense responses [77], which would be congruent with the overall absence of response of genes related to the salicylic acid pathway (Fig. 9). As genes related to cytokinin biosynthesis (*PtIPT*) and activation (*PtLOG5*) were not affected by salivary proteins of *P. passerinii*, the downregulation of cytokinin perception and signaling genes (*PtAHK4* and *PtARR2* during the assays with protoplasts and *ARR16* during the assay with *A. thaliana*) could correspond to an auxin accumulation-induced regulation loop [78, 79]. This downregulation of *PtAHK4* and *PtARR2* could also interfere with stress signaling by preventing both accumulation of jasmonate and activation of *PtPR1* [80] (Fig. 9).

**Figure 9:**
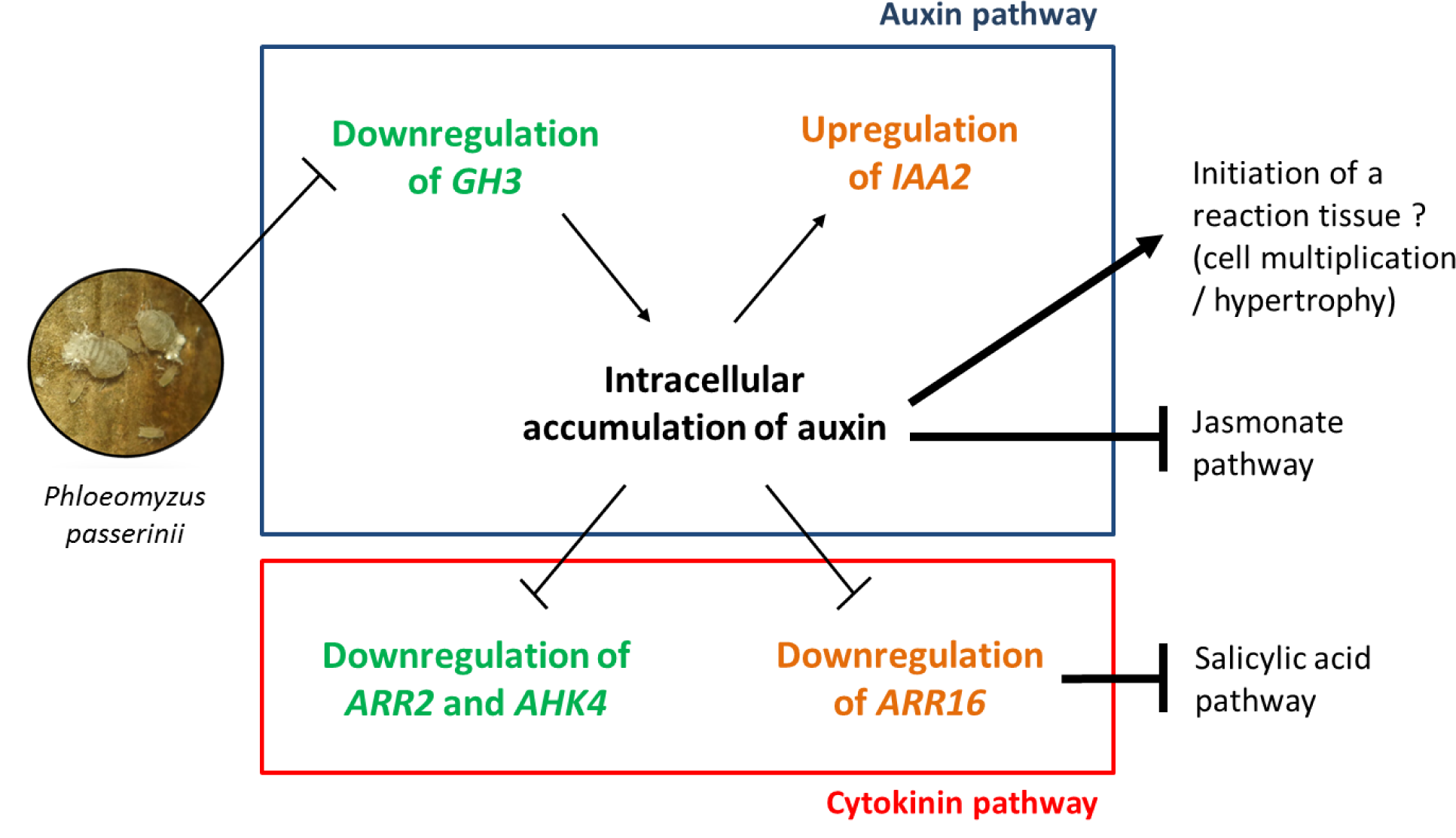
Graphical summary of effects of salivary extracts of *Phloeomyzus passerinii* on gene expression in auxin and cytokinin pathways during a compatible interaction based on observations with *in vivo* RT-qPCR assays with protoplasts (bold green) and heterologous *in planta* assay (bold orange).

The intracellular accumulation of active auxin forms could also contribute to the targeted cell hypertrophy and multiplication commonly observed in the reaction tissues induced by *P. passerinii* in susceptible poplar genotypes [20] (Fig. 9). Auxin accumulation, as a result of a reduction in both *GH3* activity and auxin transport, is also probably involved in the initiation and development of root galls by cyst and root-knot nematodes [81]. However, exposure of protoplasts of the susceptible genotype to the salivary extracts of *P. passerinii* did not affect expression of genes involved in the cell-division cycle, or weakly downregulated them. Only one gene involved in cell-division cycle (i.e. *PtCDK20*) was differentially expressed during interactions with either a susceptible or a resistant host genotype. Additional experiments considering genes coding for cell wall remodeling enzymes, activated early during root-knot nematode-induced giant cell differentiation, or for cytoskeleton should provide further insight into the mechanisms associated with the differentiation of hypertrophied cells in this system [73].

In conclusion, our transcriptomic analysis of the saliva of *P. passerinii* and *M. persicae* showed that *P. passerinii* probably secretes a highly peculiar saliva, filled with potential effectors that may interfere with several plant secondary messengers and signaling pathways. Our *in vivo* approach with protoplasts and *in planta* approach with a heterologous *A. thaliana*-system confirmed the ability of salivary extracts of *P. passerinii* to interfere with host response during interactions with a susceptible host genotype. As expected auxin and cytokinin pathways were affected, probably to impair biotic stress signaling but also to reconfigure host metabolism and anatomy. Although the saliva of *P. passerinii* and *M. persicae* were very different, interactions with non-host species and with a resistant genotype of a host species led to quite similar host responses, with a different intensity however, and few differences in biotic stress signaling and cytokinin metabolism. Additional modalities including different populations of *P. passerinii*, different poplar genotypes with intermediate resistance levels [18], and additional host metabolic pathways could be considered in future experiments. For instance, investigating how genes coding for NB-LRR proteins, involved in the induction of effector-triggered immunity following the recognition of parasite effectors [10, 82], respond to salivary extracts of *P. passerinii* should give further insights into the molecular processes underpinning failed and successful host infestation by *P. passerinii*. Similarly, investigating the response of genes coding for early nodulin-like proteins, recently characterized in the genome of *P. trichocarpa* [83], could help to unravel the molecular mechanisms underpinning host manipulation, since these genes are targeted by several gall-inducing organisms [84, 85].

## Acknowledgements

We thank Léa Fléchon for her assistance with transcriptome analyses. We are also grateful to the reviewers for their valuable comments.

## Appendices

**Figure S1:**
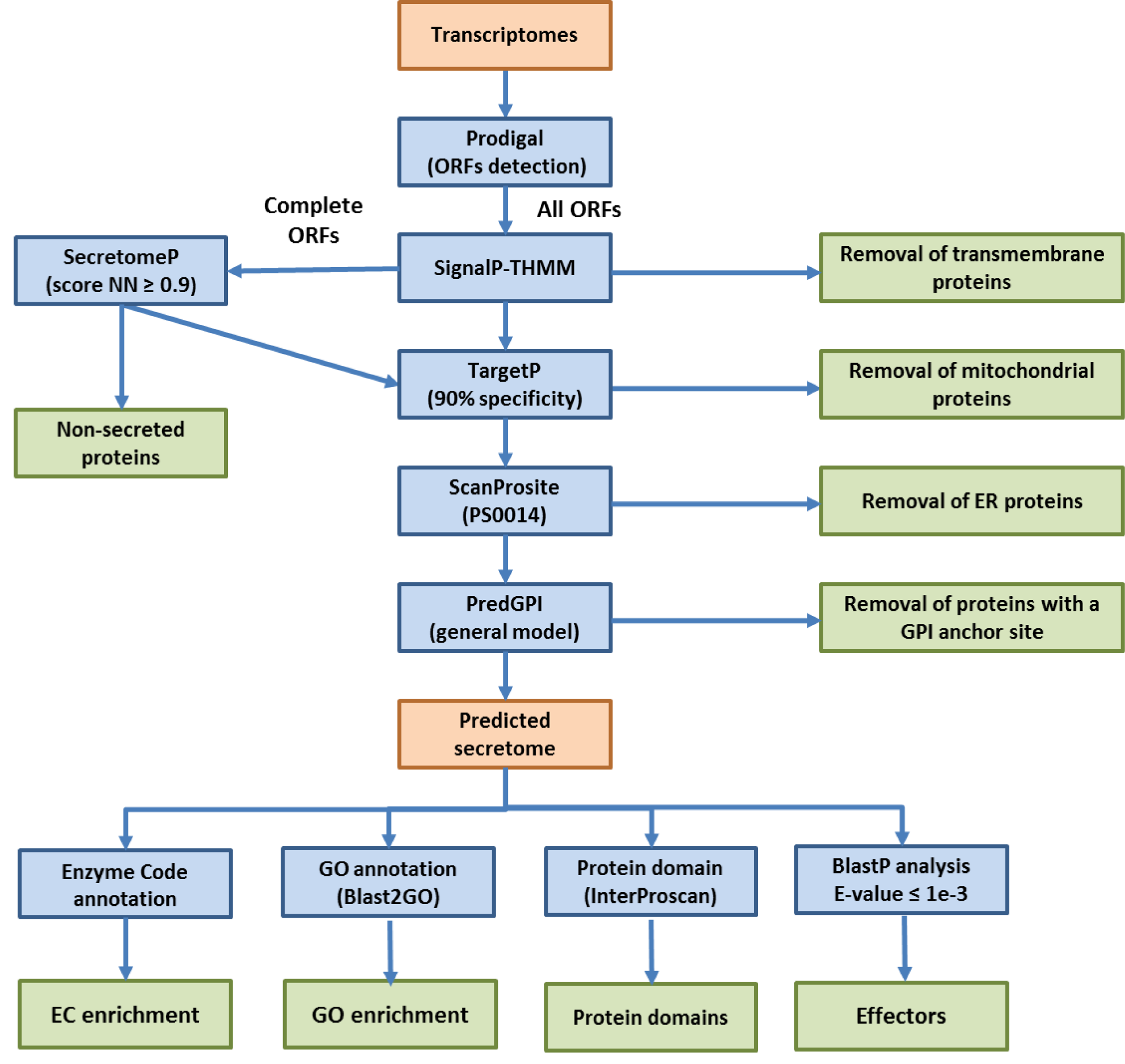
Pipeline for the annotation process. Data are in orange, analyses in blue, results in green.

**Fig. S2.**
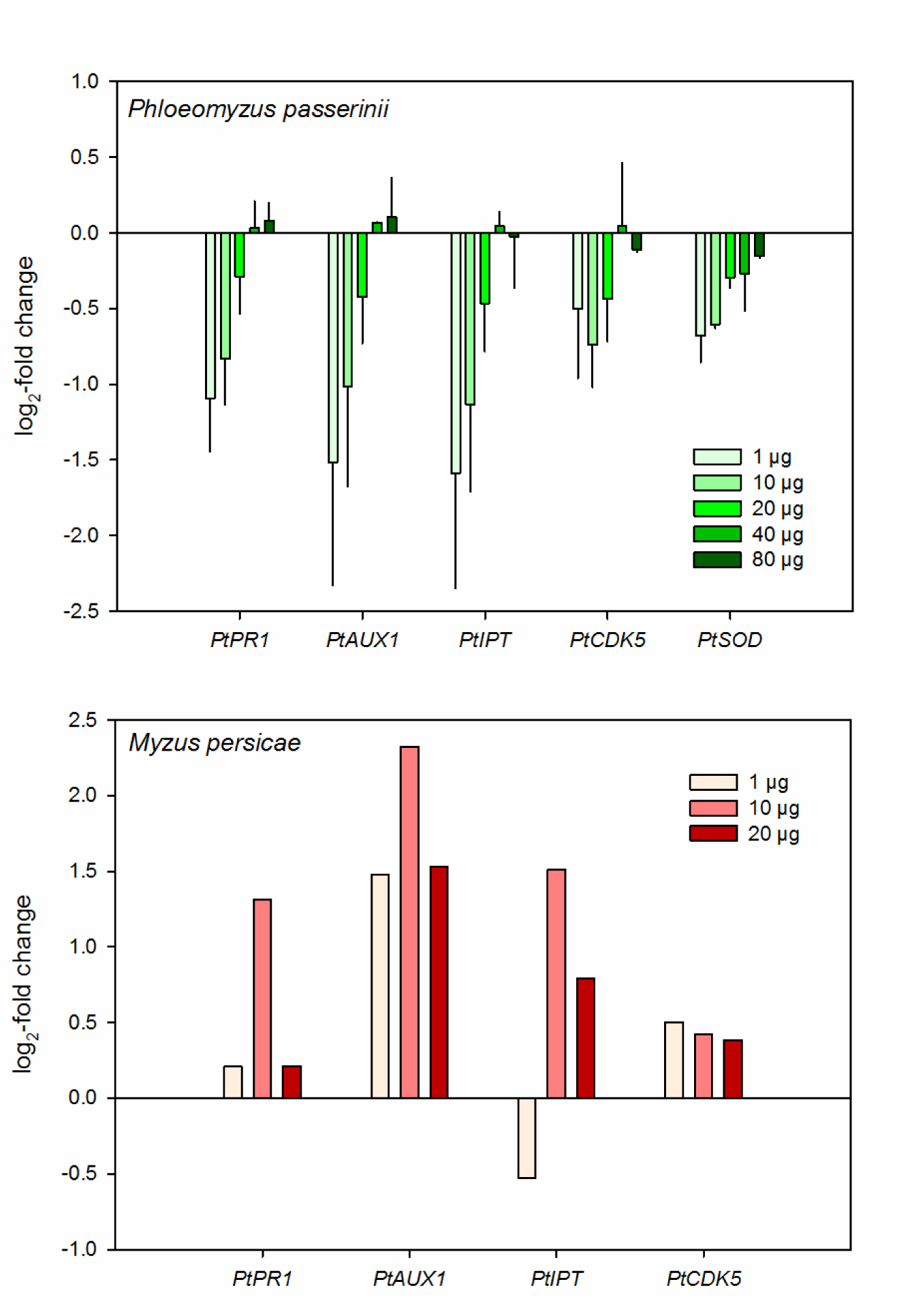
Mean (± SE) log_2_-fold changes of several poplar genes after incubation of I-214 poplar protoplasts with different doses (1, 10, 20, 40 and 80 µg) of salivary proteins of *Phloeomyzus passerinii* (upper graph) and *Myzus persicae* (lower graph). Two biological replicates were performed for *P. passerinii* and one for *M. persicae*.

**Fig. S3.**
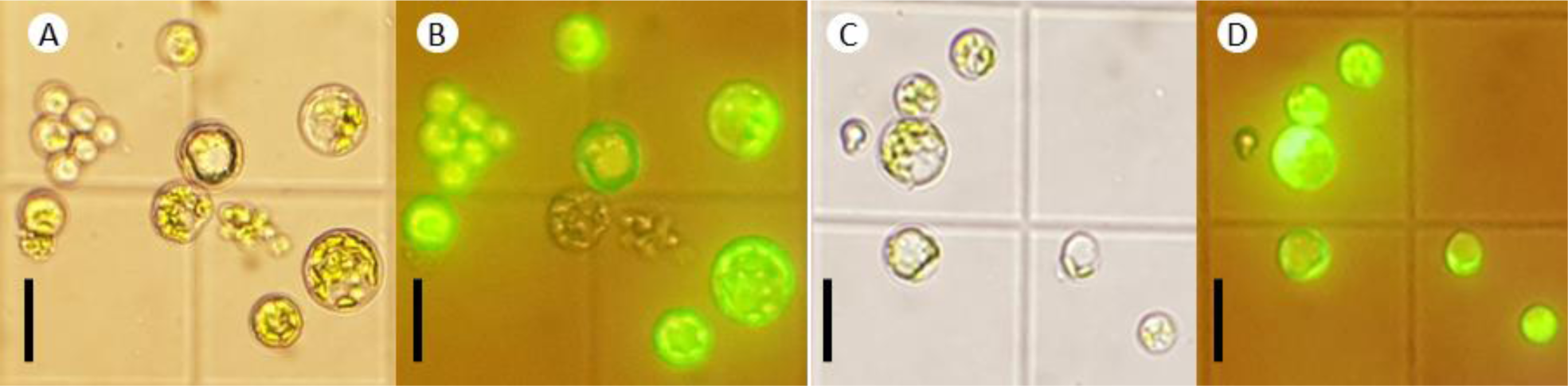
FDA assay on poplar protoplasts after 3 h of incubation with salivary proteins of *Phloeomyzus passerinii* (A, B) and *Myzus persicae* (C, D). Black bars represent 25 µm. Most protoplasts were fluorescent after treatment, indicating that incubation with salivary proteins of aphids did not affect their survival.

**Table S1:** Illumina MiSeq sequencing and assembly statistics for *Phloeomyzus passerinii* salivary transcriptome.

|  |  |
| --- | --- |
| Number of raw read-pairs | 15,453,942 |
| Number of read-pairs after quality check* | 11,578,269 |
| Number of assembled transcripts | 36,312 |
| Average transcript length (bases) | 858 |
| N50 (bases) | 1273 |
| Transcriptome completeness (%)** | 86.0 complete + 4.4 partial |

\* the reads were quality-clipped before performing the assembly with Trimmomatic v0.32. (Bolger *et al.* 2014).
\*\* assessed on 1066 core genes

**Table S2:**
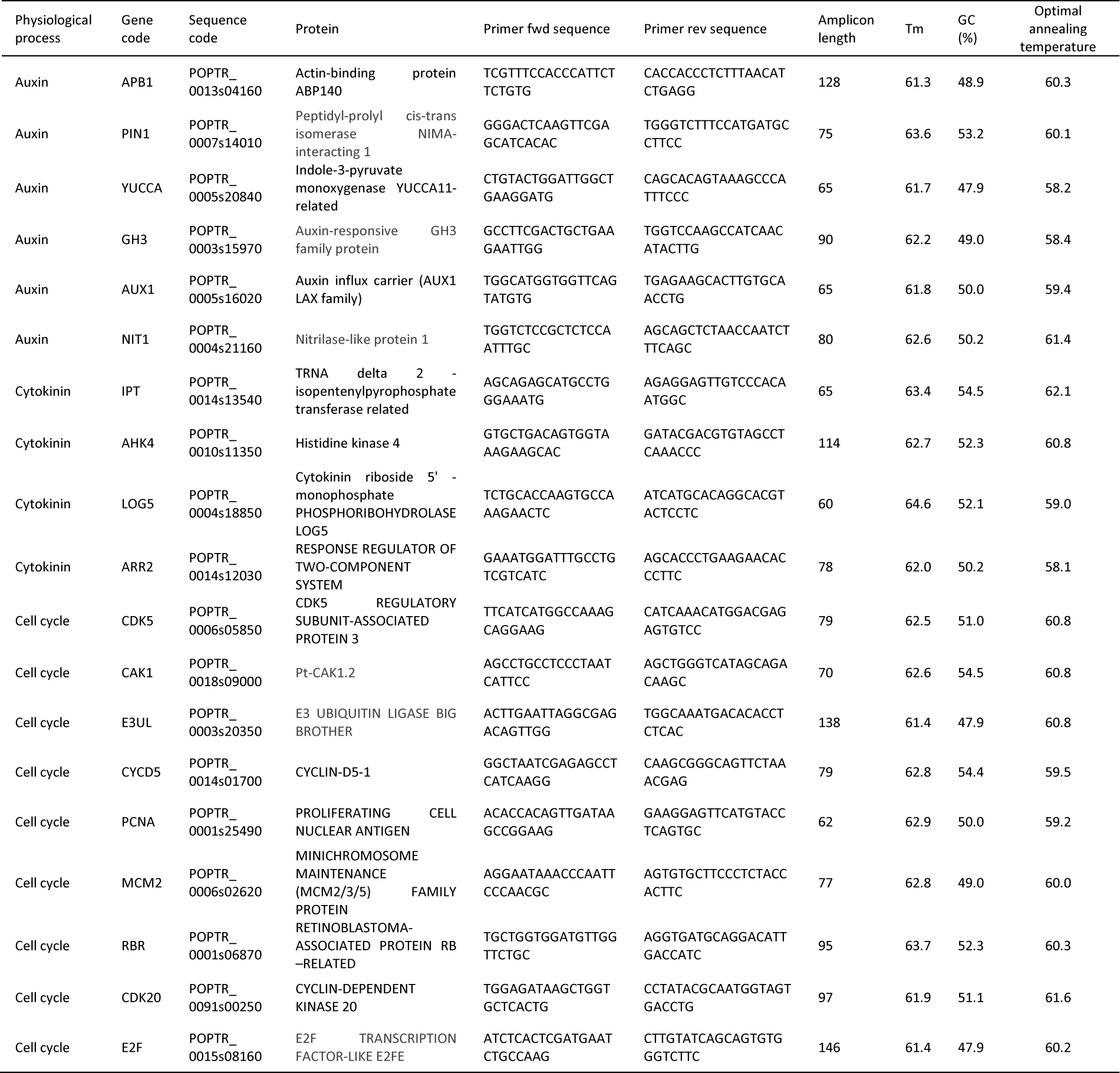
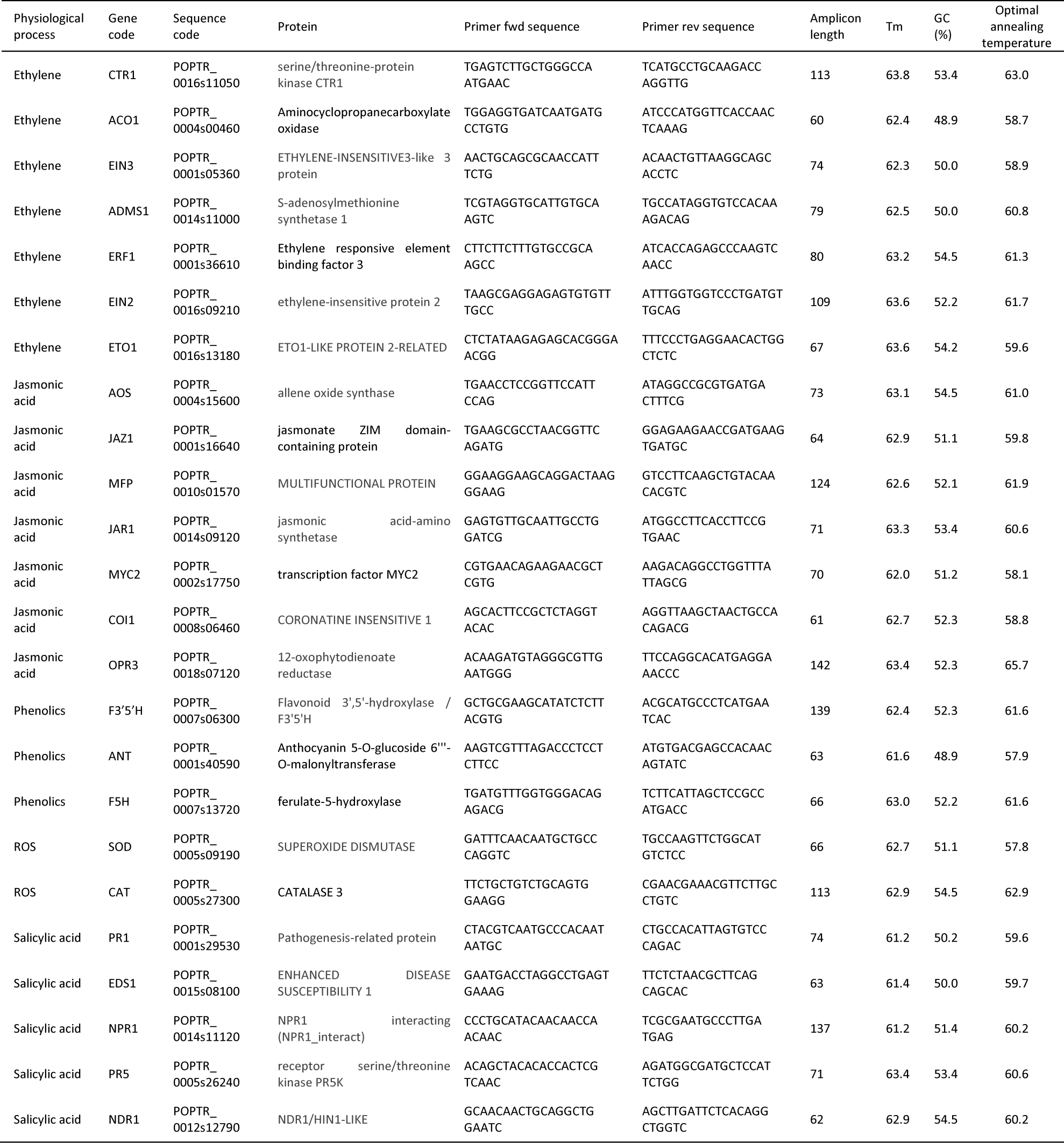
Genes considered for RT-qPCR, their sequence code in Phytozome and primers sequences.

**Table S3:**
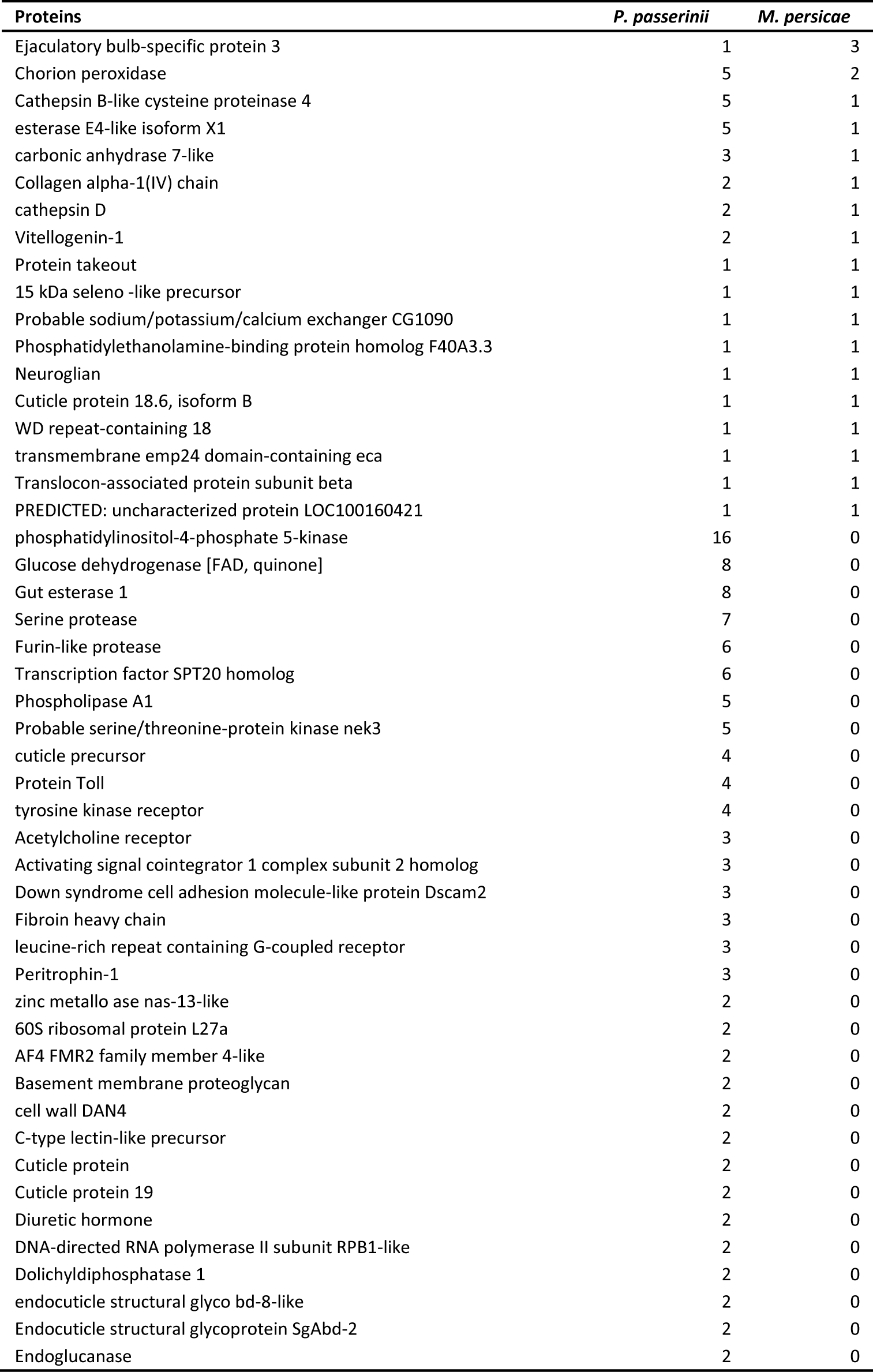
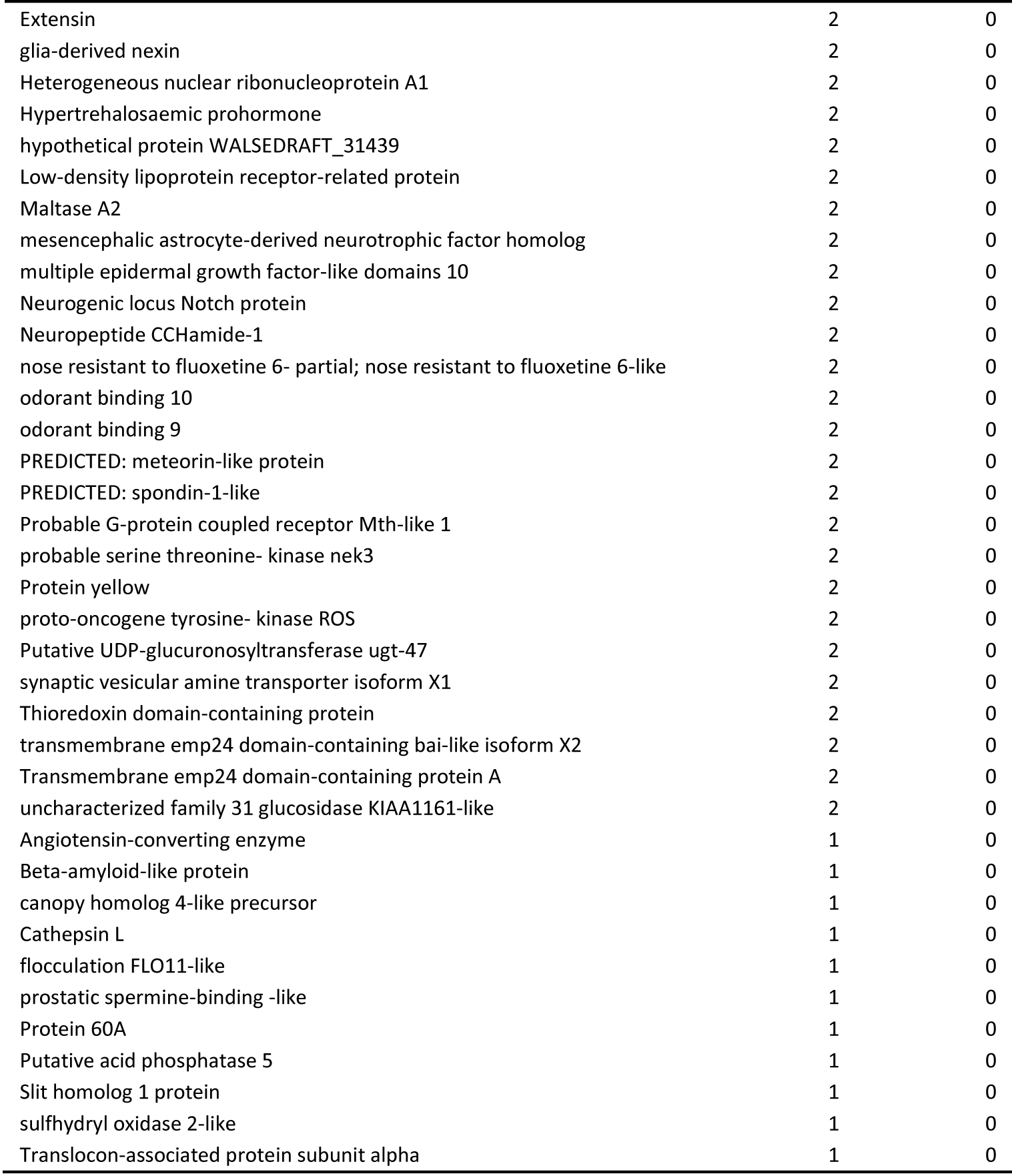
Number of orthologous proteins between the salivary transcriptomes of *Phloeomyzus passerinii* and *Myzus persicae*.

**Table S4:** Gene ontology annotations of genes from salivary glands of *Phloeomyzus passerinii*. All genes belonged to the Molecular Function category.

| Gene Ontology (GO) | Number of sequences |
| --- | --- |
| protein binding | 55 |
| nucleic acid binding | 19 |
| structural constituent of cuticle | 16 |
| phosphatidylinositol phosphate kinase activity | 15 |
| serine-type endopeptidase activity | 15 |
| zinc ion binding | 13 |
| chitin binding | 13 |
| DNA binding | 13 |
| metal ion binding | 12 |
| calcium ion binding | 12 |
| odorant binding | 9 |
| ATP binding | 9 |
| hydrolase activity | 7 |
| protein dimerization activity | 7 |
| hormone activity | 6 |
| oxidoreductase activity | 6 |
| cysteine-type endopeptidase activity | 6 |
| peroxidase activity | 5 |
| metalloendopeptidase activity | 5 |
| nucleotide binding | 5 |
| heme binding | 5 |
| carboxylic ester hydrolase activity | 4 |
| DNA-binding transcription factor activity | 4 |
| peptide-methionine (S)-S-oxide reductase activity | 4 |
| protein-hormone receptor activity | 3 |
| triglyceride lipase activity | 3 |
| nuclease activity | 3 |
| RNA binding | 3 |
| N-acetyltransferase activity | 3 |
| catalytic activity | 3 |
| transporter activity | 3 |
| acetylcholine-gated cation-selective channel activity | 3 |
| transferase activity, transferring hexosyl groups | 3 |
| hydrolase activity, hydrolyzing O-glycosyl compounds | 3 |
| peptidase activity | 3 |
| chitinase activity | 3 |
| metallopeptidase activity | 3 |
| aspartic-type endopeptidase activity | 2 |
| 3'-5' exonuclease activity | 2 |
| serine-type endopeptidase inhibitor activity | 2 |
| hydrolase activity, acting on glycosyl bonds | 2 |
| oxidoreductase activity, acting on CH-OH group of donors | 2 |
| cellulase activity | 2 |
| leucine-tRNA ligase activity | 2 |
| structural constituent of cell wall | 2 |
| acid phosphatase activity | 2 |
| G-protein coupled receptor activity | 2 |
| heparin binding | 2 |
| scavenger receptor activity | 2 |
| sulfuric ester hydrolase activity | 2 |
| cation binding | 2 |
| protein tyrosine kinase activity | 2 |
| growth factor activity | 2 |
| transferase activity | 2 |
| extracellular matrix structural constituent | 2 |
| transition metal ion binding | 2 |
| carbohydrate binding | 2 |
| obsolete signal transducer activity | 2 |
| cholinesterase activity | 2 |
| RNA-DNA hybrid ribonuclease activity | 2 |
| iron ion binding | 2 |
| aminoacyl-tRNA editing activity | 2 |
| serine-type peptidase activity | 2 |
| GTP binding | 2 |
| transferase activity, transferring glycosyl groups | 2 |
| flavin adenine dinucleotide binding | 2 |
| structural constituent of ribosome | 2 |
| G-protein beta/gamma-subunit complex binding | 1 |
| cysteine-type peptidase activity | 1 |
| mRNA binding | 1 |
| DNA-directed 5'-3' RNA polymerase activity | 1 |
| transmembrane receptor protein serine/threonine kinase activity | 1 |
| lysozyme activity | 1 |
| unfolded protein binding | 1 |
| omega peptidase activity | 1 |
| iron-sulfur cluster binding | 1 |
| alpha,alpha-trehalase activity | 1 |
| protein kinase binding | 1 |
| sequence-specific DNA binding | 1 |
| voltage-gated potassium channel activity | 1 |
| endopeptidase inhibitor activity | 1 |
| transmembrane transporter activity | 1 |
| protein kinase activity | 1 |
| copper ion binding | 1 |
| DNA-directed DNA polymerase activity | 1 |
| endonuclease activity | 1 |
| carboxypeptidase activity | 1 |
| O-acyltransferase activity | 1 |
| hedgehog receptor activity | 1 |
| isomerase activity | 1 |
| proton transmembrane transporter activity | 1 |
| GTPase activator activity | 1 |
| cis-stilbene-oxide hydrolase activity | 1 |
| carbonate dehydratase activity | 1 |
| metallocarboxypeptidase activity | 1 |
| cation transmembrane transporter activity | 1 |
| phosphatase activity | 1 |
| extracellular ligand-gated ion channel activity | 1 |
| cysteine dioxygenase activity | 1 |
| ion channel inhibitor activity | 1 |
| transmembrane receptor protein tyrosine kinase activity | 1 |
| protein heterodimerization activity | 1 |
| polysaccharide binding | 1 |
| G-protein coupled receptor binding | 1 |
| 1-alkyl-2-acetyl-glycerophosphocholine esterase activity | 1 |
| sulfotransferase activity | 1 |
| ATPase activity, coupled to transmembrane movement of substances | 1 |
| ubiquitin protein ligase binding | 1 |
| phospholipase A2 activity | 1 |
| poly(A)-specific ribonuclease activity | 1 |
| obsolete signal transducer, downstream of receptor, with serine/threonine kinase activity | 1 |
| glucosylceramidase activity | 1 |
| hydrolase activity, acting on carbon-nitrogen (but not peptide) bonds | 1 |
| ionotropic glutamate receptor activity | 1 |
| signaling receptor binding | 1 |
| GPI-anchor transamidase activity | 1 |
| extracellularly glutamate-gated ion channel activity | 1 |
| dolichyl-diphosphooligosaccharide-protein glycotransferase activity | 1 |
| thiol oxidase activity | 1 |
| phosphatidylinositol N-acetylglucosaminyltransferase activity | 1 |
| ion channel activity | 1 |
| oxidoreductase activity, acting on the aldehyde or oxo group of donors, NAD or NADP as acceptor | 1 |
| peptidyl-dipeptidase activity | 1 |
| FMN binding | 1 |
| electron transfer activity | 1 |
| lipid transporter activity | 1 |
| transferase activity, transferring acyl groups other than amino-acyl groups | 1 |
| beta-glucuronidase activity | 1 |
| alpha-mannosidase activity | 1 |
| GTPase activity | 1 |
| hydrolase activity, acting on carbon-nitrogen (but not peptide) bonds, in linear amides | 1 |
| protein serine/threonine kinase activity | 1 |
| succinate dehydrogenase activity | 1 |
| mannosyl-oligosaccharide 1,2-alpha-mannosidase activity | 1 |

## References

[1] S.A. Hogenhout, R.A. Van der Hoorn, R. Terauchi, S. Kamoun, Emerging concepts in effector biology of plant-associated organisms. Mol. Plant-Microbe Int., 22 (2009), 115–122.

[2] S.A. Hogenhout, J.I. Bos, Effector proteins that modulate plant–insect interactions. Curr. Opin. Plant Biol., 14 (2011), 422–428.

[3] J. Win, A. Chaparro-Garcia, K. Belhaj, D.G.O. Saunders, K. Yoshida, S. Dong, S. Schornack, C. Zipfel, S.A. Hogenhout, S. Kamoun, Effector biology of plant-associated organisms: concepts and perspectives. Cold Spring Harb. Symp. Quant. Biol. 77 (2012), 235–247.

[4] T. Will, W.F. Tjallingii, A. Thönnessen, A.J. van Bel. Molecular sabotage of plant defense by aphid saliva. Proc. Nat. Acad. Sci., 104 (2007), 10536–10541.

[5] A. Guiguet, G. Dubreuil, M.O. Harris, H. Appel, J.C. Schultz, M.H. Pereira, D. Giron, Shared weapons of blood- and plant-feeding insects: surprising commonalities for manipulating hosts. J. Ins. Physiol., 84 (2016), 4–21.

[6] D. Giron, E. Huguet, G.N. Stone, M. Body, Insect-induced effects on plants and possible effectors used by galling and leaf-mining insects to manipulate their host-plant. J. Ins. Physiol., 84 (2016), 70–89.

[7] L.Q. Chen, SWEET sugar transporters for phloem transport and pathogen nutrition. New Phytol., 201 (2014), 1150–1155.

[8] J. Sandström, A. Telang, N.A. Moran, Nutritional enhancement of host plants by aphids - a comparison of three aphid species on grasses. J. Ins. Physiol., 46 (2000), 33–40.

[9] M.G. Mitchum, R.S. Hussey, T.J. Baum, X. Wang, A.A. Elling, M. Wubben, E.L. Davis, Nematode effector proteins: an emerging paradigm of parasitism. New Phytol., 199 (2013), 879–894.

[10] P.N. Dodds, J.P. Rathjen, Plant immunity: towards an integrated view of plant–pathogen interactions. Nat. Rev. Genet., 11 (2010), 539.

[11] L. Lo Presti, D. Lanver, G. Schweizer, S. Tanaka, L. Liang, M. Tollot, A. Zuccaro, S. Reissmann, R. Kahmann, Fungal effectors and plant susceptibility. Ann. Rev. Plant Biol., 66 (2015), 513–545.

[12] M. Franceschetti, A. Maqbool, M.J. Jiménez-Dalmaroni, H.G. Pennington, S. Kamoun, M.J. Banfield, Effectors of filamentous plant pathogens: commonalities amid diversity. Microbiol. Mol. Biol. Rev., 81 (2017), e00066–16.

[13] I. Kaloshian, L.L. Walling, Hemipteran and dipteran pests: effectors and plant host immune regulators. J. Integr. Plant Biol., 58 (2016), 350–361.

[14] S. Basu, S. Varsani, J. Louis, Altering plant defenses: herbivore-associated molecular patterns and effector arsenal of chewing herbivores. Mol. Plant-Microbe Int., 31(2017), 13–21.

[15] J.J. Stuart, M.S. Chen, R. Shukle, M.O. Harris, Gall midges (Hessian flies) as plant pathogens. Ann. Rev. Phytopathol., 50 (2012), 339–357.

[16] J. Stuart Insect effectors and gene-for-gene interactions with host plants. Curr. Opin. Ins. Sci., 9 (2015), 56–61.

[17] D. Giron, G. Dubreuil, A. Bennett, F. Dedeine, M. Dicke, L.A. Dyer, M. Erb, M.O. Harris, E. Huguet, I. Kaloshian, A. Kawakita, C. Lopez-Vaamonde, T.M. Palmer, T. Petanidou, M. Poulsen, A. Sallé, J.-C. Simon, J. Terblanche, D. Thiery, N.K. Whiteman, H.A. Woods, S. Pincebourde (2018). Promises and challenges in insect–plant interactions. Entomol. Exp. Appl., 166 (2017), 319–343.

[18] A. Sallé, S. Pointeau, S. Bankhead-Dronnet, C. Bastien, F. Lieutier, Unraveling the tripartite interactions among the woolly poplar aphid, its host tree, and their environment: a lead to improve the management of a major tree plantation pest? Ann. For. Sci., 74 (2017), 79.

[19] S. Pointeau, A. Ameline, F. Laurans, A. Sallé, Y. Rahbé, S. Bankhead-Dronnet, F. Lieutier, Exceptional plant penetration and feeding upon cortical parenchyma cells by the woolly poplar aphid. J. Ins. Physiol., 58 (2012), 857–866.

[20] F. Dardeau, E. Deprost, F. Laurans, V. Lainé, F. Lieutier, A. Sallé, Resistant poplar genotypes inhibit pseudogall formation by the wooly poplar aphid, *Phloeomyzus passerinii* Sign., Trees, 28 (2014), 1007–1019.

[21] A. Sallé, R. Jerger, C. Vincent-Barbaroux, O. Baubet, D. Dahuron, S. Bourgerie, F. Lieutier, Tree-killing aphid dramatically reduces bark contents in carbohydrates and nitrogen compounds. For. Ecol. Manage., 407 (2018), 23–30.

[22] F. Dardeau, M. Body, A. Berthier, F. Miard, J. P. Christidès, M. Feinard-Duranceau, F. Brignolas, D. Giron, F. Lieutier, A. Sallé, Effects of fertilisation on amino acid mobilisation by a plant-manipulating insect. Ecol. Entomol., 40 (2015), 814–822.

[23] F. Dardeau, S. Pointeau, A. Ameline, F. Laurans, A. Cherqui, F. Lieutier, A. Sallé, Host manipulation by a herbivore optimizes its feeding behaviour. Anim. Behav., 95 (2014), 49–56.

[24] J.F. Tooker, A.M. Helms, Phytohormone dynamics associated with gall insects, and their potential role in the evolution of the gall-inducing habit. J. Chem. Ecol., 40 (2014), 742–753.

[25] N. Harmel, E. Létocart, A. Cherqui, P. Giordanengo, G. Mazzucchelli, F. Guillonneau, E. De Pauw, E. Haubruge, F. Francis, Identification of aphid salivary proteins: a proteomic investigation of *Myzus persicae*. Ins. Mol. Biol., 17 (2008), 165–174.

[26] M. De Vos, G. Jander, *Myzus persicae* (green peach aphid) salivary components induce defence responses in *Arabidopsis thaliana*. Plant Cell Environ., 32 (2009), 1548–1560.

[27] Z. Ren, A.J. Harris, R.B. Dikow, E. Ma, Y. Zhong, J. Wen, Another look at the phylogenetic relationships and intercontinental biogeography of eastern Asian–North American Rhus gall aphids (Hemiptera: Aphididae: Eriosomatinae): Evidence from mitogenome sequences via genome skimming. Mol. Phylogenet. Evol., 117 (2017), 102–110.

[28] A. Bishopp, S. El-Showk, D. Weijers, B. Scheres, J. Friml, E. Benková, A. Pekka Mähönen, Y. Helariutta, A mutually inhibitory interaction between auxin and cytokinin specifies vascular pattern in roots. Curr. Biol., 21 (2011), 917–926.

[29] T. Kiba, H. Yamada, T. Mizuno, Characterization of the ARR15 and ARR16 response regulators with special reference to the cytokinin signaling pathway mediated by the AHK4 histidine kinase in roots of *Arabidopsis thaliana*. Plant Cell Physiol., 43 (2002), 1059–1066.

[30] D. Zerbino, E. Birney, Velvet: algorithms for de novo short read assembly using de Bruijn graphs. Gen. Res., (2008), gr-074492.

[31] M.H. Schulz, D.R. Zerbino, M. Vingron, E. Birney, Oases: robust *de novo* RNA-seq assembly across the dynamic range of expression levels. Bioinformatics, 28 (2012), 1086–1092.

[32] Y. Yang, S.A. Smith, Optimizing *de novo* assembly of short-read RNA-seq data for phylogenomics. BMC Genomics, 14 (2013), 328.

[33] O. Nishimura, Y. Hara, S. Kuraku S. gVolante for standardizing completeness assessment of genome and transcriptome assemblies. Bioinformatics, 33 (2017), 3635–3637.

[34] D. Hyatt, G.L. Chen, P.F. LoCascio, M.L. Land, F.W. Larimer, L.J. Hauser, Prodigal: prokaryotic gene recognition and translation initiation site identification. BMC Bioinformat., 11 (2010), 119.

[35] T.N. Petersen, S. Brunak, G. von Heijne, H. Nielsen, SignalP 4.0: discriminating signal peptides from transmembrane regions, Nat. Methods, 8 (2011), 785.

[36] R. Ji, H. Yu, Q. Fu, H. Chen, W. Ye, S. Li, Y. Lou, Comparative transcriptome analysis of salivary glands of two populations of rice brown planthopper, *Nilaparvata lugens*, that differ in virulence. PloS One, 8 (2013), e79612.

[37] J.D. Bendtsen, L.J. Jensen, N. Blom, G. von Heijne, S. Brunak, Feature-based prediction of non-classical and leaderless protein secretion. Prot. Eng. Des. Sel., 17 (2004), 349–356.

[38] O. Emanuelsson, H. Nielsen, S. Brunak, G. von Heijne, Predicting subcellular localization of proteins based on their N-terminal amino acid sequence. J. Mol. Biol., 300 (2000), 1005–1016.

[39] A. Pierleoni, P.L. Martelli, R. Casadio, PredGPI: a GPI-anchor predictor. BMC Bioinformat., 9 (2008), 392.

[40] A. Conesa, S. Götz, J.M. García-Gómez, J. Terol, M. Talón, M. Robles, Blast2GO: a universal tool for annotation, visualization and analysis in functional genomics research. Bioinformat., 21 (2005), 3674– 3676.

[41] P. Jones, D. Binns, H.Y. Chang, M. Fraser, W. Li, C. McAnulla, H. McWilliam, J. Maslen, A. Mitchell, G. Nuka, S. Pesseat, A.F. Quinn, A. Sangrador-Vegas, M. Scheremetjew, S.-Y. Yong, R. Lopez, S. Hunter, InterProScan 5: genome-scale protein function classification. Bioinformat., 30 (2014), 1236–1240.

[42] Y. Wang, D. Coleman-Derr, G. Chen, Y.Q. Gu. OrthoVenn: a web server for genome wide comparison and annotation of orthologous clusters across multiple species. Nucleic Acids Res., 43 (2015), W78–W84.

[43] S. Pundir, M.J. Martin, C. O’Donovan, Uniprot protein knowledgebase, in: C. Wu, C. Arighi, K. Ross (Eds). Protein Bioinformatics. Methods in Molecular Biology, vol 1558. Humana Press, New York, 2017, 41–55.

[44] K. Guo, W. Wang, L. Luo, J. Chen, Y. Guo, F. Cui, Characterization of an aphid-specific, cysteine-rich protein enriched in salivary glands. Biophys. Chem., 189 (2014), 25–32.

[45] M. Jaouannet, P.A. Rodriguez, P. Thorpe, C.J.G. Lenoir, R. MacLeod, C. Escudero-Martinez, J.I.B. Bos, Plant immunity in plant–aphid interactions. Front. Plant Sci., 5 (2014), 663.

[46] Y. Pan, J. Zhu, L. Luo, L. Kang, F. Cui, High expression of a unique aphid protein in the salivary glands of *Acyrthosiphon pisum*. Physiol. Mol. Plant Pathol., 92 (2015), 175–180.

[47] D.A. Elzinga, G. Jander, The role of protein effectors in plant–aphid interactions. Curr. Opin. Plant Biol., 16 (2013), 451–456.

[48] A. Cherqui, W.F. Tjallingii, W.F. Salivary proteins of aphids, a pilot study on identification, separation and immunolocalisation. J. Ins. Physiol., 46 (2000), 1177–1186.

[49] J.L. Yang, R. Yang, A.C. Cheng, R.Y. Jia, M.S. Wang, S.H. Zhang, Five-minute purification of PCR products by new-freeze-squeeze method. J. Food Agric. Environ., 8 (2010), 32–33.

[50] M.E. Maffei, A. Mithöfer, W. Boland, Insects feeding on plants: rapid signals and responses preceding the induction of phytochemical release. Phytochemistry, 68 (2007), 2946–2959.

[51] F.H. Wu, S.C. Shen, L.Y. Lee, S.H. Lee, M.T. Chan, C.S. Lin, Tape-Arabidopsis Sandwich-a simpler Arabidopsis protoplast isolation method. Plant Meth., 5 (2009), 16.

[52] J. Guo, J.L. Morrell-Falvey, J.L. Labbé, W. Muchero, U.C. Kalluri, G.A. Tuskan, J.G. Chen, Highly efficient isolation of Populus mesophyll protoplasts and its application in transient expression assays. PloS One, 7 (2012), e44908.

[53] S. Arvidsson, M. Kwasniewski, D.M. Riaño-Pachón, B. Mueller-Roeber, QuantPrime--a flexible tool for reliable high-throughput primer design for quantitative PCR. BMC Bioinformat., 9 (2008), 465.

[54] Z. Tong, Z. Gao, F. Wang, J. Zhou, Z. Zhang, Selection of reliable reference genes for gene expression studies in peach using real-time PCR. BMC Mol. Biol., 10 (2009), 71.

[55] J.E. Malamy, P.N. Benfey, Organization and cell differentiation in lateral roots of *Arabidopsis thaliana*. Development, 124 (1997), 33–44.

[56] R Core Team (2013). R: A language and environment for statistical computing. R Foundation for Statistical Computing, Vienna, Austria. URL http://www.R-project.org/.

[57] R. Suzuki, H. Shimodaira, Pvclust: an R package for assessing the uncertainty in hierarchical clustering. Bioinformatics, 22 (2006), 1540–1542.

[58] Y. Cuesta-Astroz, F.S. de Oliveira, L.A. Nahum, G. Oliveira, Helminth secretomes reflect different lifestyles and parasitized hosts. Int. J. Parasitol., 47 (2017), 529–544.

[59] S. Vandermoten, N. Harmel, G. Mazzucchelli, E. De Pauw, E. Haubruge, F. Francis. Comparative analyses of salivary proteins from three aphid species. Ins. Mol. Biol., 23 (2014), 67–77.

[60] W.R. Cooper, J.W. Dillwith, G.J. Puterka, Comparisons of salivary proteins from five aphid (Hemiptera: Aphididae) species. Environ. Entomol., 40 (2011), 151–156.

[61] S.J. Nicholson, S.D. Hartson, G.J. Puterka, Proteomic analysis of secreted saliva from Russian Wheat Aphid (*Diuraphis noxia* Kurd.) biotypes that differ in virulence to wheat. J. Proteomics, 75 (2012), 2252– 2268.

[62] D.A. Elzinga, G. Jander, The role of protein effectors in plant–aphid interactions. Curr. Opin. Plant Biol., 16 (2013), 451–456.

[63] S. Cambier, O. Ginis, S.J. Moreau, P. Gayral, J. Hearn, G. Stone, D. Giron, E. Huguet, J.M. Drezen, Gall wasp transcriptomes unravel potential effectors involved in molecular dialogues with oak and rose bushes. Front. Physiol., 10 (2019), 926.

[64] J.S. Ramsey, A.C.C. Wilson, M. de Vos, Q. Sun, C. Tamborindeguy, A. Winfield, G. Malloch, D.M. Smith, B. Fenton, S.M. Gray, G. Jander, Genomic resources for *Myzus persicae*: EST sequencing, SNP identification, and microarray design. BMC Genom., 8 (2007), 423–423.

[65] H. Boulain, F. Legeai, E. Guy, S. Morlière, N.E. Douglas, J. Oh, M. Murugan, M. Smith, J. Jaquiéry, J. Peccoud, F.F. White, J.C. Carolan, J.-C. Simon, A. Sugio,. Fast evolution and lineage-specific gene family expansions of aphid salivary effectors driven by interactions with host-plants. Genome Biol. Evol., 10 (2018), 1554–1572.

[66] J.C. Carolan, D. Caragea, K.T. Reardon, N.S. Mutti, N. Dittmer, K. Pappan, F. Cui, M. Castaneto, J. Poulain, C. Dossat, D. Tagu, J.C. Reese, G.R. Reeck, T.L. Wilkinson, O.R. Edwards, Predicted effector molecules in the salivary secretome of the pea aphid (*Acyrthosiphon pisum*): a dual transcriptomic/proteomic approach. J. Proteome Res., 10 (2011), 1505–1518.

[67] C. Testerink, T. Munnik, Phosphatidic acid: a multifunctional stress signaling lipid in plants. Trends Plant Sci., 10 (2005), 368–375.

[68] H.W. Xue, X. Chen, Y. Mei, Function and regulation of phospholipid signalling in plants. Biochem. J., 421 (2009), 145–156.

[69] E.O. Martinson, J.D. Hackett, C.A. Machado, A.E. Arnold, Metatranscriptome analysis of fig flowers provides insights into potential mechanisms for mutualism stability and gall induction. PloS One, 10 (2015), e0130745.

[70] I. Morkunas, V.C. Mai, B. Gabryś, Phytohormonal signaling in plant responses to aphid feeding. Acta Physiol. Plant., 33 (2011), 2057–2073.

[71] P.I. Kerchev, B. Fenton, C.H. Foyer, R.D. Hancock, Infestation of potato (*Solanum tuberosum* L.) by the peach-potato aphid (*Myzus persicae* Sulzer) alters cellular redox status and is influenced by ascorbate. Plant Cell Environ, 35 (2012), 430–440.

[72] J. Louis, J. Shah. *Arabidopsis thaliana*—*Myzus persicae* interaction: shaping the understanding of plant defense against phloem-feeding aphids. Front. Plant Sci., 4 (2013), 213.

[73] B. Favery, M. Quentin, S. Jaubert-Possamai, P. Abad. Gall-forming root-knot nematodes hijack key plant cellular functions to induce multinucleate and hypertrophied feeding cells. J. Ins. Physiol., 84 (2016), 60–69

[74] A. Mine, C. Seyfferth, B. Kracher, M.L. Berens, D. Becker, K. Tsuda, The defense phytohormone signaling network enables rapid, high-amplitude transcriptional reprogramming during effector-triggered immunity. Plant Cell, 30 (2018), 1199–1219.

[75] L. Wu, H. Chen, C. Curtis, Z.Q. Fu, Go in for the kill: How plants deploy effector-triggered immunity to combat pathogens. Virulence, 5 (2014), 710–721.

[76] K. Kazan, R. Lyons, Intervention of phytohormone pathways by pathogen effectors. Plant Cell, 26 (2014), 2285–2309.

[77] J.E. Park, J.Y. Park, Y.S. Kim, P.E. Staswick, J. Jeon, J. Yun, S.-Y. Kim, J. Kim, Y.-H. Lee, C.M. Park. GH3-mediated auxin homeostasis links growth regulation with stress adaptation response in *Arabidopsis*. J. Biol. Chem., 282 (2007), 10036–10046.

[78] B. Jones, S.A. Gunnerås, S.V. Petersson, P. Tarkowski, N. Graham, S. May, K. Dolezal, G. Sandberg, K. Ljung, Cytokinin regulation of auxin synthesis in *Arabidopsis* involves a homeostatic feedback loop regulated via auxin and cytokinin signal transduction. Plant Cell, 22 (2010), 2956–2969.

[79] G.E. Schaller, A. Bishopp, J.J. Kieber, The yin-yang of hormones: cytokinin and auxin interactions in plant development. Plant Cell, 27 (2015), 44–63.

[80] J.A. O’Brien, E. Benková, Cytokinin cross-talking during biotic and abiotic stress responses. Front Plant Sci., 4 (2013), 451.

[81] A. Karczmarek, H. Overmars, J. Helder, A. Goverse, Feeding cell development by cyst and root-knot nematodes involves a similar early, local and transient activation of a specific auxin-inducible promoter element. Mol. Plant Pathol., 5 (2004), 343–346.

[82] F.L. Goggin, Plant–aphid interactions: molecular and ecological perspectives. Curr. Opin. Plant Biol., 10 (2007), 399–408.

[83] S. Luo, W. Hu, Y. Wang, B. Liu, H. Yan, Y. Xiang, Genome-wide identification, classification, and expression of phytocyanins in *Populus trichocarpa*. Planta, 247 (2018), 1133–1148.

[84] M.-A. Cannesan, E. Nguema-Ona, Arabinogalactan proteins in root–microbe interactions. Trends Plant Sci. 18 (2013) 440–449.

[85] J. Hearn, M. Blaxter, K. Schönrogge, J.L. Nieves-Aldrey, J. Pujade-Villar, E. Huguet, J.-M. Drezen, J.D. Shorthouse, G.N. Stone, Genomic dissection of an extended phenotype: Oak galling by a cynipid gall wasp. PLoS Gen. 15 (2019).

